# PLK1 inhibition selectively kills ARID1A deficient cells through uncoupling of oxygen consumption from ATP production

**DOI:** 10.1101/2021.06.01.446664

**Authors:** Upadhyayula S. Srinivas, Gokula K. Ramachandran, Joanna D. Wardyn, Michal M. Hoppe, Yanfen Peng, Sherlly Lim, May Yin Lee, Praveen C. Peethala, Omer An, Akshay Shendre, Bryce W.Q. Tan, Tay S.C. Norbert, Patrick Jaynes, Longyu Hu, Rekha Jakhar, Karishma Sachaphibulkij, Lina H.K. Lim, Karen Crasta, Henry Yang, Patrick Tan, Dennis Kappei, Yong Wei Peng, David S.P. Tan, Shazib Pervaiz, Matteo Bordi, Silvia Campello, Wai Leong Tam, Christian Frezza, Anand D. Jeyasekharan

**Author notes:** To whom correspondence is to be addressed*: **Corresponding Author**: Dr Anand D. Jeyasekharan, Cancer Science Institute of Singapore, Centre for Translational Medicine (MD6), #13-01G, 14 Medical Drive, Singapore 117599.

## Abstract

Inhibitors of the mitotic kinase PLK1 yield objective responses in a subset of refractory cancers. However, PLK1 overexpression in cancer does not correlate with drug sensitivity, and the clinical development of PLK1 inhibitors has been hampered by the lack of patient selection marker. Using a high-throughput chemical screen, we discovered that cells deficient for the tumor suppressor ARID1A are highly sensitive to PLK1 inhibition. Interestingly this sensitivity was unrelated to canonical functions of PLK1 in mediating G2-M cell cycle transition. Instead, a whole-genome CRISPR screen revealed PLK1 inhibitor sensitivity in ARID1A deficient cells to be dependent on the mitochondrial translation machinery. We find that ARID1A knocked-out (KO) cells have an unusual mitochondrial phenotype with aberrant biogenesis, increased oxygen consumption/ expression of oxidative phosphorylation genes, but without increased ATP production. Using expansion microscopy and biochemical fractionation, we see that a subset of PLK1 localizes to the mitochondria in interphase cells. Inhibition of PLK1 in ARID1A KO cells further uncouples oxygen consumption from ATP production, with subsequent membrane depolarization and apoptosis. Knockdown of a key subunit of the mitochondrial ribosome reverses PLK1-inhibitor induced apoptosis in ARID1A deficient cells, confirming specificity of the phenotype. Together, these findings highlight a novel interphase role for PLK1 in maintaining mitochondrial fitness under metabolic stress, and a strategy for therapeutic use of PLK1 inhibitors. To translate these findings, we describe a quantitative microscopy assay for assessment of ARID1A protein loss, which could offer a novel patient selection strategy for the clinical development of PLK1 inhibitors in cancer.

**Statement of significance:** Currently, no predictive biomarkers have been identified for PLK1 inhibitors in cancer treatment. We show that ARID1A loss sensitizes cells to PLK1 inhibitors through a previously unrecognized vulnerability in mitochondrial oxygen metabolism.

## Introduction

Polo-like kinase 1 (PLK1) is a mitotic kinase which mediates G2/M cell cycle transition, centrosome maturation, microtubule-kinetochore attachment and cytokinesis(1). It is commonly overexpressed in tumors and is associated with poor survival(2), making it an attractive therapeutic target. Early phase clinical trials of PLK1 kinase inhibitors have yielded equivocal results(3,4). Several PLK1 inhibitors show clinical responses across diverse cancer types(4), but to date there are no randomized data demonstrating improvement in survival. Among current PLK1 inhibitors, Volasertib is most advanced in clinical development, having received an FDA breakthrough designation in acute myeloid leukemia(5). The combination of PLK1 inhibition with other chemotherapeutics such as gemcitabine and cytarabine improves response rates(4), but this is associated with significant toxicity. Biomarkers to select for tumors sensitive to PLK1 inhibition are therefore essential to improve the available therapeutic index: as lower doses can be combined with other agents for these cases. Interestingly, PLK1 expression itself has not proven to be a biomarker of PLK1 inhibitor efficacy(6), and the lack of a predictive biomarker has hampered efforts to refine and further the clinical use of these drugs.

Using a high-throughput chemical library screen, we identified that cells deficient for the tumor suppressor ARID1A were sensitive to PLK1 inhibitors. AT- rich interacting domain 1A (ARID1A), also called BAF250, is a subunit of the mammalian SWItch/Sucrose Non-Fermentable (SWI/SNF) or BRG1/BRM associated factors (BAF) complex. Mutations in the BAF complex are noted in 10-20% of all cancers(7), with ARID1A loss being the most frequent of these. It behaves as a tumor suppressor, with its loss promoting tumor differentiation and progression(8,9). The BAF complex is a family of chromatin remodeling proteins capable of using ATP to alter nucleosomes and thereby chromatin accessibility(10). Multiple subunits of the BAF complex mediate the interaction of the complex with DNA and histones, culminating in ATP-dependent chromatin modulation by the core ATPases Brahma (BRM) and BRM/SWI2 related gene 1 (BRG1)(11). By modulating access to DNA, ARID1A contributes to the maintenance of genomic stability by affecting recruitment of proteins such as ATR and TOP2A(9) to sites of DNA damage(12,13). However, ARID1A’s tumor suppressor role is also ascribed to the regulation of a variety of genes through its enrichment at promoters and enhancers and mediation of RNAPII pause-release(14). A drastic change in the metabolism of cells upon ARID1A loss is considered to be an important tumorigenic effect, and the SWI/SNF complex was indeed originally described in the setting of cellular metabolism(15).

In our study, we discovered that the rewiring of metabolism in ARID1A-deficient cells leads to an altered mitochondrial phenotype with aberrant biogenesis, increased oxygen consumption and expression of oxidative phosphorylation genes, but without increased ATP production. We describe a novel localization of PLK1 to the mitochondria in interphase. Inhibition of PLK1 in ARID1A deficient cells exacerbates mitochondrial oxygen consumption and membrane depolarization and thereby results in apoptosis. Importantly, PLK1 inhibition does not preferentially affect cell cycle progression in ARID1A deficient cells, pointing to a non-canonical cell-cycle independent role for PLK1 in maintaining mitochondrial health in cells lacking ARID1A. Overall, ARID1A loss in multiple cell types synergizes with PLK1 inhibitor activity, and thus represents a promising biomarker to refine the clinical development of Volasertib and other PLK1 inhibitors.

## Materials and Methods

### Cell culture

MCF10A cells were obtained from ATCC and cultured in DMEM/F12 supplemented with 5% horse serum, 10μg/ml insulin, 20 ng/ml epithelial growth factor, 0.5 mg/ml hydrocortisone and 100 ng/ml cholera toxin as per ATCC instructions. GES1 was a gift to Dr. Patrick Tan from from Dr. Alfred Cheng, Chinese University of Hong Kong. GES1 cells were grown in RPMI1640 supplemented with 20% FBS and 1% Pen/Strep. Cells were cultured in incubators at 37°C/ 5% CO_2_. siRNA treatment using siAURKAIP1(Horizon) was performed using Lipofectamine 2000 (Thermofisher, cat#11668500) as per the manufacturer’s instructions.

### Generation of ARID1A KO cells

ARID1A KO cells in, GES1, OVCAR3 and MCF10A were generated using CRISPR-Cas9 and validated by immunoblotting and immunofluorescence. sgRNA sequences targeting ARID1A were obtained from Broad institute sgRNA designer (FP-5′CACCGATGCATGATGCTGTCCGAC3′, RP-5′AAACGTCGGACAGCATCATGCATC 3′). Briefly, sgRNA oligos were mixed in equimolar ratios and heated to 95°C followed by cooling at room temperature overnight to form duplex oligos. Oligos were cloned into respective Cas9 plasmids and sequenced (LCV2 for OVCAR3 and GES1, FgH1tUTG for MCF10A). HEK293 cells were used to generate viruses to transduce target mammalian cells. Following transductions cells were selected with 1 μg/ml puromycin (GES1 and OVCAR3) and serial diluted to obtain single cells or single cell sorted directly (MCF10A). Serial dilution was performed to obtain single cell clones in 96 well plates. ARID1A KO clones were confirmed using immunoblotting for ARID1A and selected clones were expanded for further experiments.

### Kinase inhibitor screen and validation

Kinase library screen was performed using MCF10A-ARID1A WT and KO cells. Briefly, 1000 cells per well of each cell line were plated in 96 well plates and incubated at 37°C overnight. Next day, cells were treated with 10 μM of the kinase inhibitors from the library and incubated for 72 h at 37°C. Post incubation, cell proliferation was measured using cell titer blue assay as per manufacturer’s instructions. Fluorescence intensity was measured using Tecan infinite 200 Pro plate reader, survival fractions were plotted and Z scores calculated for the assay.

### Chemicals

Volasertib (S2235), Alisertib (MLN 8237), Dasatinib (S1021), GSK 126 (S7061) was purchased from Selleck chemicals. VE821 (SML 1415), Olaparib (AZD2461), Crystal violet (C6158) were purchased from Sigma-Aldrich. Cisplatin (CAY13119) was obtained from Cayman chemicals. JC-1 (Cat#T3168), Mitotracker red, were obtained from ThermofisherScientific. Cell titer blue was purchased from Promega (G8081).

### Western blotting and antibodies

Whole cell lysates were prepared in RIPA lysis buffer (Thermofisher, cat#89900) supplemented with cOmplete™ Protease Inhibitor Cocktail tablets (Sigma-Aldrich, cat#11697498001). 7 or 10 % SDS-PAGE gels were employed for analysis. Antibodies used are detailed in supplementary table1.

### Cell proliferation assay (Cell Titer Blue)

For cell proliferation assays, 1000 cells per well were plated in a 96 well plate and incubated overnight at 37°C. Next day, cells were treated with indicated concentrations of chemotherapeutics for 72 h. After incubation Cell Titer Blue was directly added to the wells and incubated for 4h at 37 °C followed by fluorescence measurement as per manufacturer’s instructions. The cell survival percentage was calculated as a fraction of DMSO controls and plotted in PRISM™.

### Cell cycle analysis

Cells were plated at a density of 2×10^5^ cells per well of a 6 well plate and treated with Volasertib (0-100 nM) for 24 h. After incubation, cells were harvested, fixed in ice cold ethanol. Cells were then washed with PBS and incubated with RNase A for 30 m at 37°C followed by PI staining at room temperature for 10 m. Data acquisition was done using BD LSRII and analysis was performed using FlowJo™ V10.

### Phospho-histone H3 staining

Cells were plated at a density of 2×10^5^ cells per well of a 6 well plate, treated with Volasertib (0, 10 and 100 nM) for 24 h. Post treatment, cells were fixed with 70% ice cold ethanol, washed with PBS, permeabilized with 0.25% Triton-X 100 in PBS for 10 m at room temperature, and blocked with 3% BSA in PBS for 30 m at room temperature. Samples were incubated with Anti-phospho H3 antibody (1:50, CST#9701) overnight at 4°C, followed by 3x washes with PBS and incubation with Alexa 488 (1:500, Thermofisher, A11094) secondary antibody for 1 h at room temperature and 3x washes with PBS. 50 μg/ml PI with RNAseA was added to the samples and incubated at room temperature for 30 m in dark. Samples were analyzed using BD LSRII and data analyzed with FlowJo ™V10.

### Annexin staining

Cells were plated at a density of 2×10^5^ per well of a 6 well plate, treated with Volasertib (0, 10 and 100 nM) for 24 h. Cells were collected and washed 3x with PBS followed by washing with Annexin binding buffer. Cells were incubated with Annexin V for 30 m at 37°C followed by incubation with PI for 10 m at RT. Data was acquired using BD LSRII and analyzed with FlowJo ™V10.

### Quantitative immunofluorescence assays

Cells were grown at a density of 8000 per well in a 96 well plate and treated with DMSO, 10 nM Volasertib, and 10 μM Cisplatin for 24 h. Samples were fixed with 4% PFA, washed with PBS, permeabilized with 0.25% Triton-X-100, blocked with 5% BSA in PBS for 30 m at RT and incubated in primary antibodies at 4°C overnight (γ-H2AX 1:1000). Samples were washed 3xPBS and incubated with fluorescently labeled Alexa-fluor secondary antibodies and Hoechst 33342 (ThermoFisher Scientific, H3570) to stain DNA for 45 m at RT in dark. Quantitative fluorescence imaging was performed with Operetta high-content imaging system (PerkinElmer) and signal quantified using manufacturer’s algorithms.

### Mitochondrial membrane potential determination with JC-1

Cells were plated at a density of 2×10^5^ cells per well of a 6 well plate, treated with Volasertib (0, 10 and nM) for 24 h. Post treatment, 200,000 cells per condition were collected, washed with PBS and incubated with dye JC-1 for 30 m at 37°C. Samples were analyzed using BD LSRII and data analyzed with FlowJo™ V10.

### Seahorse assay

Cells were plated at a density of 2×10^4^ cells per well in a seahorse XF96 (Agilent) cell culture microplates and cultured overnight at 37°C. Sensor cartridge was hydrated overnight in a non-CO_2_ incubator and cells were treated with 10 nM volasertib for 8 h. Assay media was prepared by supplementing seahorse RPMI with 1 mM pyruvate, 2mM glutamine and 10 mM glucose. Working stocks of oligomycin (1.5 μM), FCCP (0.5 μM) and Rotenone/antimycin A (0.5 μM) were prepared and loaded into respective ports of sensor cartridge. The machine was calibrated with calibration plate and sensor cartridge and replaced with cell culture microplate when ready. Simultaneously, another 96 well plate was used for normalization with DAPI staining. Results were analyzed using WAVE software and graphs plotted in PRISM™.

### Confocal microscopy

Cells were grown at a density of 4×10^4^ on coverslips, treated with chemotherapeutics for indicated times, fixed with 4% PFA for 10 m at RT, permeabilized with 0.5% triton-X-100 for 15 m at RT, blocked with 3% BSA for 30 m at RT and stained with antibodies against PLK1 (208G4), alpha-tubulin (ab7291), ATP5B (ab 14730), Tom20 (SC-17764) at 4°C overnight. Next day, cells were washed 3X in blocking solution and incubated with fluorescently labelled secondary antibody. Cells were washed three times with PBS and DNA was stained with Hoechst 33342. Zeiss LSM 880 was used to capture images.

### Quantitative polymerase chain reaction

Cells were plated in a 6 well plate at a density of 2×10^5^ per well and RNA was isolated using QIAGEN RNeasy mini kit as per manufacturer’s instructions. RNA concentration was estimated by nanodrop and 1 μg of RNA was used to synthesize cDNA with iScript cDNA synthesis kit™.(Biorad 1708890) qPCR was performed using Precision™ 2X qPCR master mix (Primerdesign) following manufacturer’s protocol. Primer sequences used are provided in supplementary table 2.

## Results

### ARID1A loss synergizes with PLK1 inhibition in-vitro and in-vivo

ARID1A mutations in cancers are typically associated with loss of protein function or/and expression(16). We generated ARID1A knockout (KO) MCF10A (non-transformed breast epithelial) cells using CRISPR-Cas9 (Fig. 1A). We then performed a high-throughput chemical library screen with FDA-approved kinase inhibitors. Of the 60 clinical-grade kinase inhibitors tested, MCF10A-ARID1A KO cells were exquisitely sensitive to PLK1 inhibition by TAK-960 (Fig. 1B). To validate this primary screen, we confirmed the observation using Volasertib, a potent FDA-approved PLK1 inhibitor. Volasertib preferentially killed ARID1A KO cells in comparison to WT cells across a range of doses (Fig. 1C). To rule out effects arising from cell line context specificity, we generated ARID1A KO cells in another cellular model – a gastric epithelial cell line GES1 (Fig. 1D) – as ARID1A loss is prevalent in gastric cancer(17). Cell proliferation assays (Fig. 1E) and colony formation assays confirmed the sensitization of GES1-ARID1A KO cells to Volasertib (Fig. 1F and S1B). Similarly, cell proliferation measured with crystal violet staining showed reduced proliferation of ARID1A KO cells in the presence of Volasertib (Fig. S1A). The differential sensitivity of ARID1A KO cells to PLK1 inhibition was higher than that with other small molecule inhibitors previously published to be synthetically lethal with ARID1A loss (8,13,18)(Fig. S1C, S1D and S1E). To demonstrate if ARID1A loss constitutes a therapeutic vulnerability for Volasertib, we setup an in-vivo experiment to compare the growth of ARID1A WT and KO tumor xenografts in the presence or absence of Volasertib administration. As MCF10A and GES1 are immortalized epithelial cell lines which are non-tumorigenic we generated ARID1A KO in OVCAR3 (Ovarian cancer) cells (Fig. S1F), another cancer type where ARID1A loss is common. Similar to MCF10A and GES1, OVCAR3-ARID1A KO cells were sensitive to Volasertib in-vitro (Fig. S1G and S1H). OVCAR3-ARID1A WT/KO cells were injected subcutaneously into the flanks of 5 mice per group. Tumors developed 10 days after injection, and the animals were treated with 15mg/kg Volasertib once per week with a control group given equivalent vehicle (V/V) (Fig. 1G). Both the volasertib treated and control group maintained healthy body mass measured twice per week (Fig. S1I) indicating that the dosage of drug did not have any major side effect on the mice. In accordance with our in-vitro studies, OVCAR3-ARID1A KO tumors showed a greater reduction in tumor size after Volasertib treatment than WT tumors (Fig. 1H and 1I). Together, these experiments suggest that PLK1 inhibition may constitute a therapeutic strategy for ARID1A mutant cancers.

**Fig. 1:**
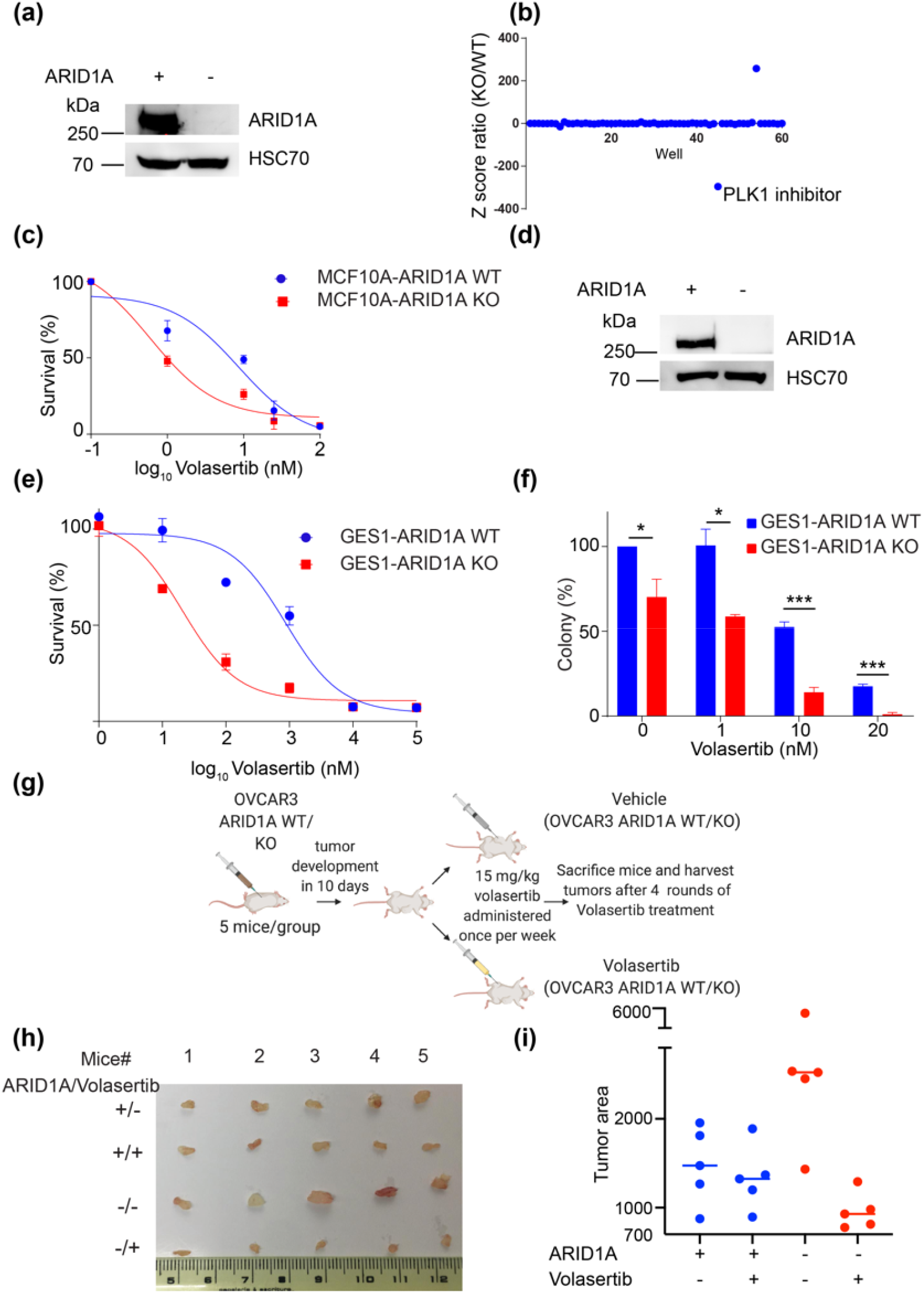
ARID1A loss induces cell-cycle independent sensitivity to PLK1 inhibitors. a) Western blot of ARID1A in MCF10A-ARID1A WT and KO cells generated using CRISPR-Cas9. HSC70 is used as a loading control, b) Dot plot representing Z-score ratios of each drug from the screen performed using an FDA-approved kinase inhibitor library, demonstrating decreased viability of KO cells in the presence of a PLK1 inhibitor. As the ARID1A WT and KO cells were screened in different plates, we use the Z-score ratio to compare them (ratio of Z-scores of a given drug for ARID1A KO/WT is calculated as observed value for a given drug-plate mean/ plate Standard Deviation), c) Cell proliferation assay confirming synergy between Volasertib and loss of ARID1A. MCF10A-ARID1A WT and KO cells were plated in 96 well plates and treated with Volasertib for 72 h, with cell viability measured using Cell Titer Blue. Experiments were performed with 4 technical replicates for each concentration of Volasertib and DMSO controls. Error bars represent SEM from three independent biological experiments, d) Western blot of ARID1A in GES1-ARID1A WT and KO cells generated using CRISPR-Cas9. HSC70 is used as a loading control, e) Cell proliferation assay confirming synergy between Volasertib and GES1-ARID1A KO cells. GES1-ARID1A WT and KO cells were plated in 96 well plates and treated with Volasertib for 72 h, with cell viability measured using Cell Titer Blue, f) Quantitation of colonies from colony formation assay experiments (n=3). Error bars represent SEM, p values of *<0.05, ***<0.001, Student’s t-test, for WT vs KO cells. Cells were plated at a dilution of 200 cells/well in a 6 well plate and treated with different concentrations of Volasertib for 10 days. Surviving cells were stained with crystal violet and the single colonies were counted, g) Schematic representation of mouse xenograft experiments. Nude (SCID) mice received sub-cutaneous injections of OVCAR3-ARID1A WT or KO cells. Tumors were visible 10 days’ post injection, after which the animals were treated with Volasertib (15mg/kg body weight) or vehicle once per week for 4 weeks. Body mass and tumor volumes were recorded twice per week. Mice were sacrificed at the end of 4 weeks of treatment and tumors were isolated and imaged, h) Images of OVCAR3-ARID1A WT/KO tumors isolated from mice after sacrifice, i) Dot plot of tumor area from h) showing changes in tumor area in OVCAR3-ARID1A KO vs OVCAR3-ARID1A KO + volasertib treatment. Tumor area was calculated from the images in image J using the ruler to set scale and applying free hand tool to measure the tumor area.

### PLK1 inhibition does not preferentially impair cell cycle arrest or progression in ARID1A deficient cells

As PLK1 activity is essential for progression through the cell cycle, we checked whether the sensitivity of ARID1A KO cells to Volasertib was explained by differences in cell cycle progression. Both GES1-ARID1A WT and KO isogenic cell lines showed similar baseline cell cycle profiles (Fig. 2A and 2B). Addition of Volasertib led to a concentration-dependent G2/M arrest, but this was similar in ARID1A WT and KO cells (Fig. 2A and 2B). PLK1 is particularly critical for the G2/M transition. We therefore investigated if PLK1 inhibition in the context of ARID1A loss altered entry into mitosis possibly leading to increased mitotic catastrophe. Since G2/M measurement by 4N DNA content does not discriminate between G2 arrested cells and cells that have entered mitosis, we measured phospho-histone H3-a marker for cells that have entered mitosis. pHH3 staining by flow cytometry was similar in ARID1A WT and KO cells. Volasertib treatment progressively increased pHH3 positive cells as expected, but no significant changes were noted between the isogenic cell lines (Fig. 2C and 2D). Together, these results suggest that PLK1 inhibition does not cause increased cell death in ARID1A deficient cells through cell-cycle related effects.

**Fig. 2:**
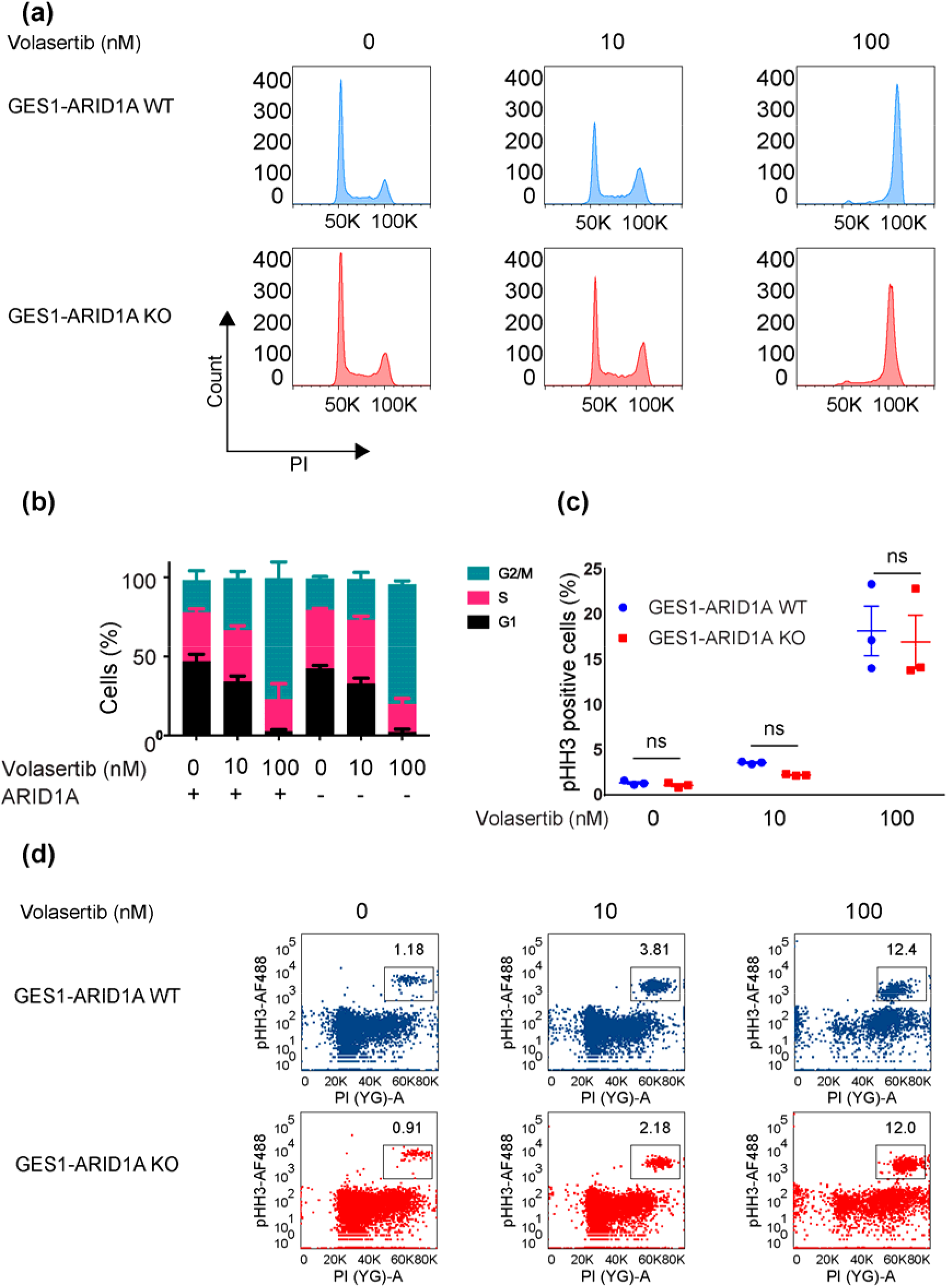
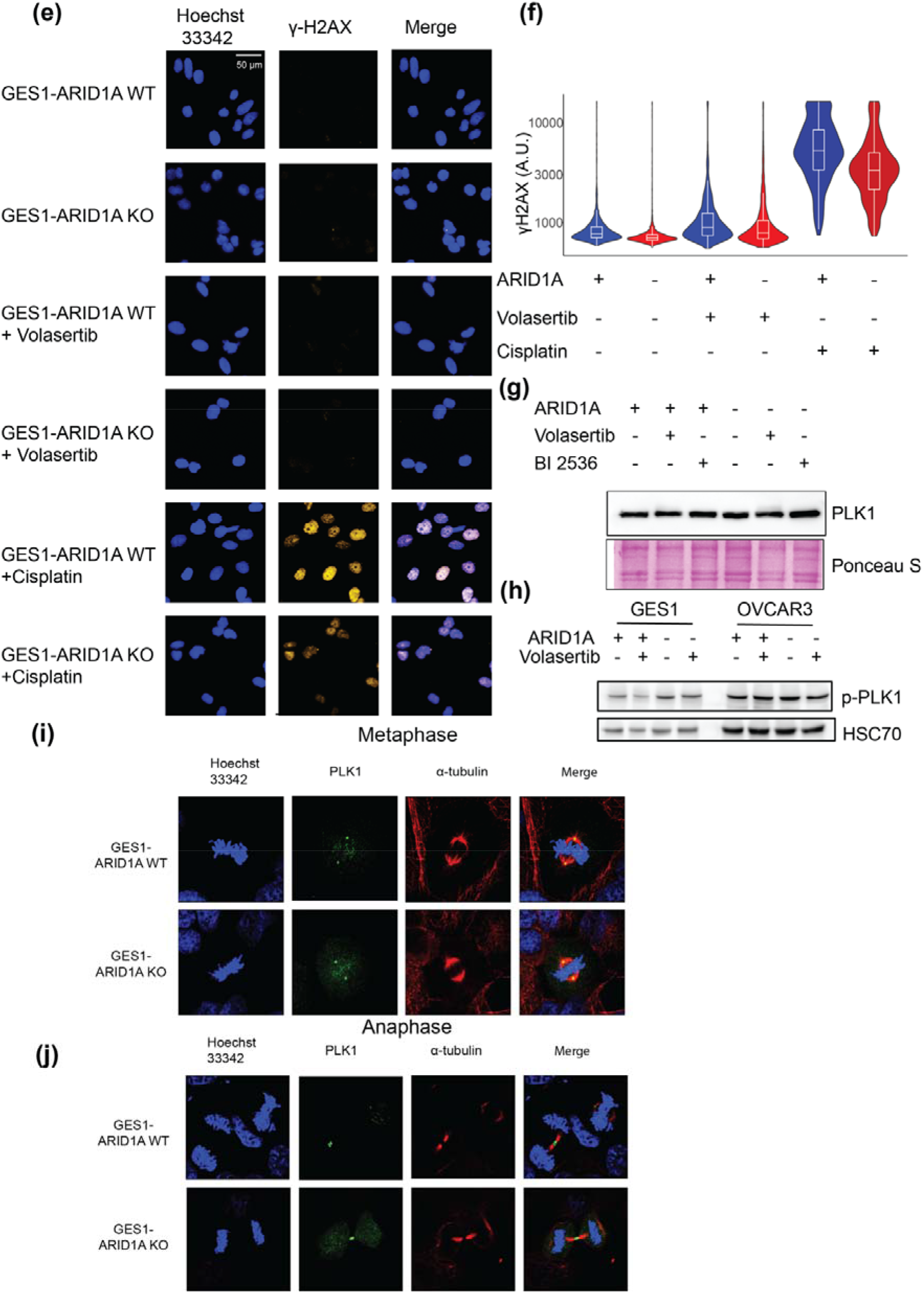
ARID1A loss does not affect cell cycle profiles, DDR activation and PLK1 levels after PLK1 inhibition. a) Cell cycle distribution histograms of GES1-ARID1A WT (blue) and GES1-ARID1A KO (red) measured by propidium iodide (PI) staining using flow cytometry. Cells were plated in a 6 well plate and treated with Volasertib for 24/48 h. Post treatment cells were fixed in ethanol and stained with PI. Measurements were done using BD FACS and histograms obtained using FlowJo TM V10, b) Bar-charts represent cells in G1 phase (black), S phase (red), G2/M phase (green). No statistical significance was observed between GES1-ARID1A WT and KO cells with identical treatments. Quantification of cell cycle phases was done using FlowJo TM V10. Error bars represent SEM of 3 independent biological experiments, Student’s t-test, c) Dot plot showing quantitation of pHH3 positive cells in GES1-ARID1A WT and GES1-ARID1A KO cells from three independent biological experiments. Statistical significance was analyzed using one-way ANOVA. d) Representative images of pHH3 staining measured by flow cytometry in GES1-ARID1A WT and GES1-ARID1A KO cells treated with Volasertib. Cells were fixed in ethanol post treatment with Volasertib for 24 h. Cells were permeabilized with 0.5% triton-x-100, blocked in 5% BSA, incubated with pHH3 antibody overnight at 4oC followed by secondary antibody incubation for 30 m at RT. Measurements were done using BD FACS. e) Representative fluorescence images of GES1-ARID1A WT and GES1-ARID1A KO cells treated with DMSO, Volasertib, Cisplatin and stained for γ-H2AX, obtained on an Operetta high-throughput microscope. Approximately 1000 cells were imaged per well, and replicates of 3 wells were imaged per condition, f) Violin wrapped-around boxplot showing frequency distribution of γ-H2AX intensity in GES1-ARID1A WT and GES1-ARID1A KO cells and treatment with DMSO, 10 nM Volasertib and 10 μM Cisplatin. Cells were plated at a density of 8000/well in a 96 well plate with three technical replicates for each treatment. Cisplatin treatment serves as a positive control for the γ-H2AX antibody. Data obtained on an Operetta high-throughput microscope. g) Western blot of PLK1 in GES1-ARID1A WT and GES1-ARID1A KO cells treated with PLK1 inhibitors Volasertib/BI2536 or DMSO showing no change in total protein levels of PLK1 between the ARID1A isogenic cell lines. Cells were cultured in 6 well dishes treated with 10 nM Volasertib/BI2536 and harvested 24 h later in RIPA buffer. Samples were prepared for SDS PAGE and run on 10% SDS-PAGE gels. Post transfer, the membrane was probed for PLK1 (CST #208G4) and PonceauS was used as a general loading control. h) Western blot analysis of phospho-PLK1 (T-210) in GES1/OVCAR3 ARID1A WT and KO cells treated with Volasertib showing no difference in phospho-PLK1 levels. HSC70 was used as a loading control. i) Cells were grown on coverslips, fixed with 4% formaldehyde and permeabilized with 0.5% triton-X-100. Following permeabilization, cells were blocked in 5% BSA/PBS for 30 m at RT and then stained with alpha tubulin/PLK1 O/N at 4°C. The cells were washed with 5% BSA/PBS thrice and incubated with secondary antibody for 45 m at RT. Imaging was performed using Carl Zeiss 880 confocal microscope. Confocal images of GES1-ARID1A WT and KO cells in Metaphase showing no difference in PLK1 localization between the isogenic cell lines. Alpha tubulin was used to stain microtubules. ⍰-tubulin (red) and PLK1 (green). Scale bar = 10 μm j) The same as i) but for Anaphase, showing no difference in PLK1 localization between the isogenic cell lines. Alpha tubulin was used to stain microtubules. IZ-tubulin (red) and PLK1 (green). Scale bar = 10 μm

Activation of the DNA damage response (DDR) is a common upstream signal for initiating apoptosis(19). We sought to determine whether differences in DDR activation could explain the increased apoptosis observed in ARID1A-deficient cells, particularly since ARID1A loss has been shown to influence the response to replication stress(13). We compared DDR activation in ARID1A WT and KO cells by quantitative immunofluorescence analysis for γ-H2AX, with and without the addition of Volasertib, as PLK1 facilitates the recovery of cells from the G2/M checkpoint after DNA damage(20). There was no significant difference for γ-H2AX between ARID1A WT and KO cells, even upon exposure to Volasertib (Fig. 2E and 2F). In contrast, the use of cisplatin as a positive control for DDR activation, led to a significant increase in γ-H2AX in both WT and KO cells. The extent of γ-H2AX after cisplatin treatment was relatively lower for the KO cells, in keeping with an established role for ARID1A in the sensing of replication stress (8) (Fig. 2E and 2F). Overall, however, these data rule out the presence of baseline DDR activation in KO cells as an explanation for increased apoptotic cell death. A previous report(21) showed sensitivity of ARID1A deficient cells to inhibitors of Aurora kinase A; attributed to a direct effect of ARID1A on Aurora kinase A transcription. However, we did not see a difference in either PLK1 expression (Fig. 2G), or Aurora-A dependent phosphorylation of PLK1 (Fig. 2H) in ARID1A deficient cells suggesting that the mechanism of cell death with PLK1 inhibitors is unlikely to be a direct transcriptional regulation of either PLK1 or Aurora-A levels. Loss of ARID1A changes global gene expression due to its role as a chromatin remodeler(14). ARID1A loss could therefore indirectly alter PLK1 localization and function in mitosis, thereby accounting for PLK1 inhibitor sensitivity. However, no difference in the localization of PLK1 between WT and KO ARID1A isogenic cell lines was observed (Fig. 2I and 2J).

### Whole genome CRISPR screen identifies that mitochondrial translation is crucial for Volasertib sensitivity in ARID1A deficient cells

As the canonical functions of PLK1 were not responsible for the increased sensitivity of ARID1A KO cells to Volasertib, we performed an unbiased whole genome CRISPR screen to identify genes/ pathways that could rescue Volasertib sensitivity in ARID1A KO cells (Fig. 3A). We utilized the Brunello library for the whole genome CRISPR screen(22), which consists of 77,471 sgRNAs with an average of 4 sgRNA per gene. Our library preparation ensured almost all sgRNAs were represented normally (99.9% representation with a low GINI index of 0.05). MCF10A- ARID1A WT and KO cells transduced with the Brunello sgRNA library were selected with puromycin for 14 days prior to treatment with Volasertib. 3 weeks after continuous exposure to Volasertib, cells were harvested for genomic DNA extraction and sequencing. The sgRNA pool had a coverage of 53% for ARID1A WT and 74% for ARID1A KO and essential genes dropped out of both WT and KO cells. For further analysis we selected genes that had all 4 of their respective sgRNAs represented in both the Volasertib treated ARID1A WT and KO cells. We then ordered these selected genes according to the median of raw counts. For each gene we calculated the ratio of the medians from the Volasertib treated KO cells with respect to the Volasertib treated WT cells and plotted hits that had a ratio of more than 800 (threshold selected to include top 100 hits) (Fig. 3B). Median calculation of sgRNA raw counts preferentially enriched in ARID1A KO cells highlighted many sgRNAs targeting mitochondrial processes (Fig. 3C). Functional enrichment using g:Profiler and cytoscape also revealed mitochondrial translation as a significantly enriched pathway in ARID1A KO cells relative to WT cells (Fig. 3C and 3D). Further analysis into the sgRNAs contributing to the pathway enrichment revealed that most of the differentially enriched sgRNAs targeted mitochondrial ribosomal subunits (Fig. 3E).

**Fig. 3:**
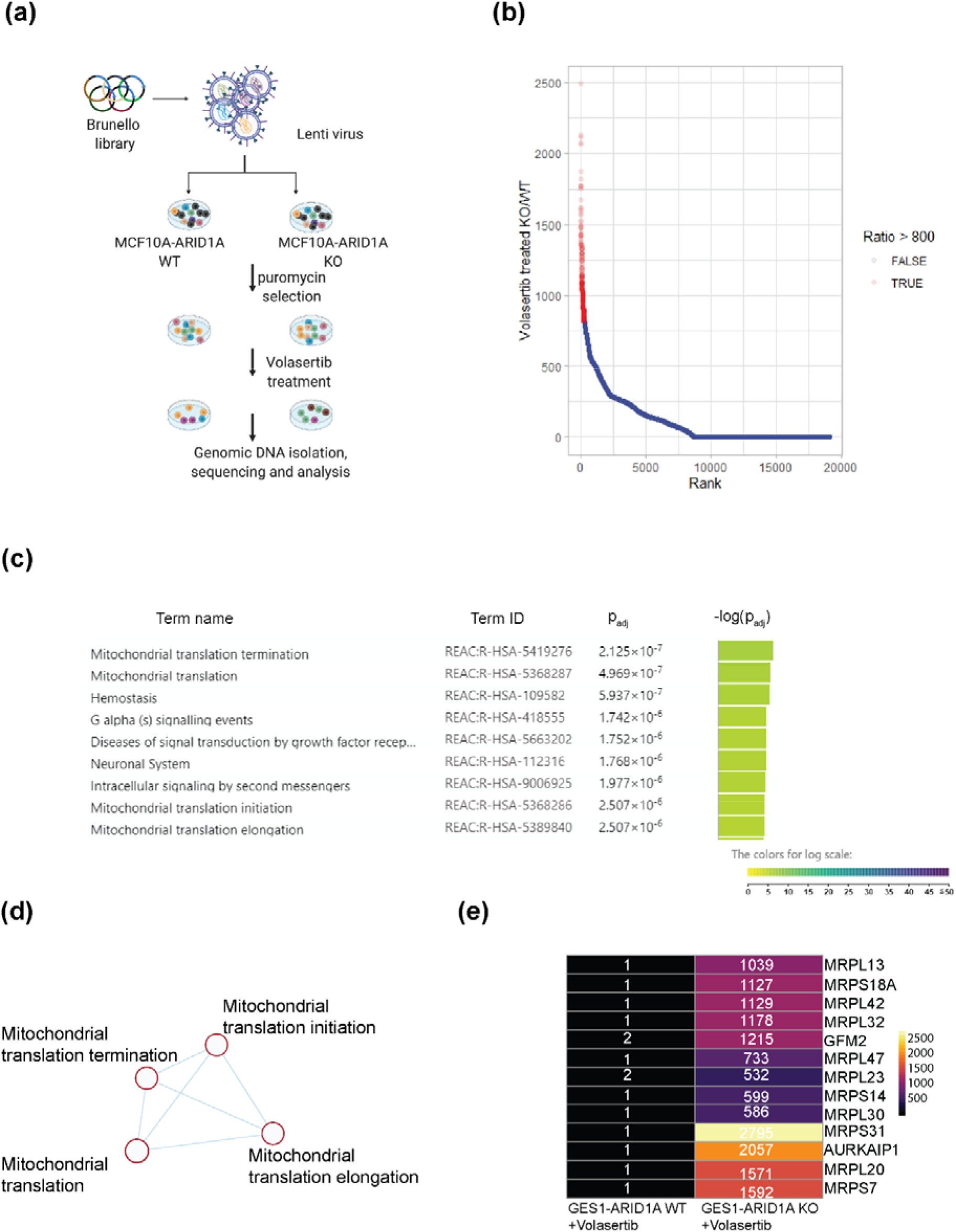
A genome-wide CRISPR screen reveals that the mitochondrial translation network is a critical determinant of Volasertib induced cell death in ARID1A KO cells. a) Schematic of the genome-wide CRISPR screen performed: the Brunello CRISPR library targeting 19,114 genes with 77,441 sgRNAs was applied via lentiviral transduction to both MCF10A-ARID1A-WT and MCF10A-ARID1A KO cells. After selection with puromycin for cells containing sgRNAs for 2 weeks, each population was treated with Volasertib for 28 days. After treatment, cells were collected and genomic DNA was extracted. Sequencing was performed to identify sgRNAs that were integrated into the genome and bioinformatic analysis followed. b) Individual sgRNAs were ranked based on the ratio of enrichment between Volasertib-treated MCF10A-ARID1A KO cells and Volasertib-treated MCF10A-ARID1A WT cells. Top enriched sgRNAs (those with a ratio greater than 800) were labelled in red. c) g:Profiler pathway analysis (https://biit.cs.ut.ee/gprofiler/gost) of top enriched sgRNAs from b) (labelled in red). Mitochondrial translation pathways were among the top enriched pathways. d) STRING analysis of mitochondrial genes obtained from c) validating the enrichment of mitochondrial translation pathways. e) Heat map demonstrating the enrichment of mitochondrial translation genes in the Volasertib-treated MCF10A-ARID1A KO cells with respect to the Volasertib-treated MCF10A-ARID1A WT cells.

### ARID1A loss leads to an aberrant mitochondrial phenotype

To extend the findings from our CRISPR screen, we evaluated the mitochondria of ARID1A WT and KO cells. High-resolution imaging of mitochondria (using immunofluorescence for Tom20) revealed clear distinctions in morphology between ARID1A WT and KO cells, with higher number of globular mitochondria in the knockout cells and reduction in long tubular (fused) mitochondria (Fig. 4A and 4B). We calculated the mitochondrial spread using the mitofootprint algorithm from MiNA image j plugin(23) to confirm that mitochondria are less spread-out in ARID1A KO cells compared to WT cells (Fig. 4C, 4D, and 4E).

**Fig. 4:**
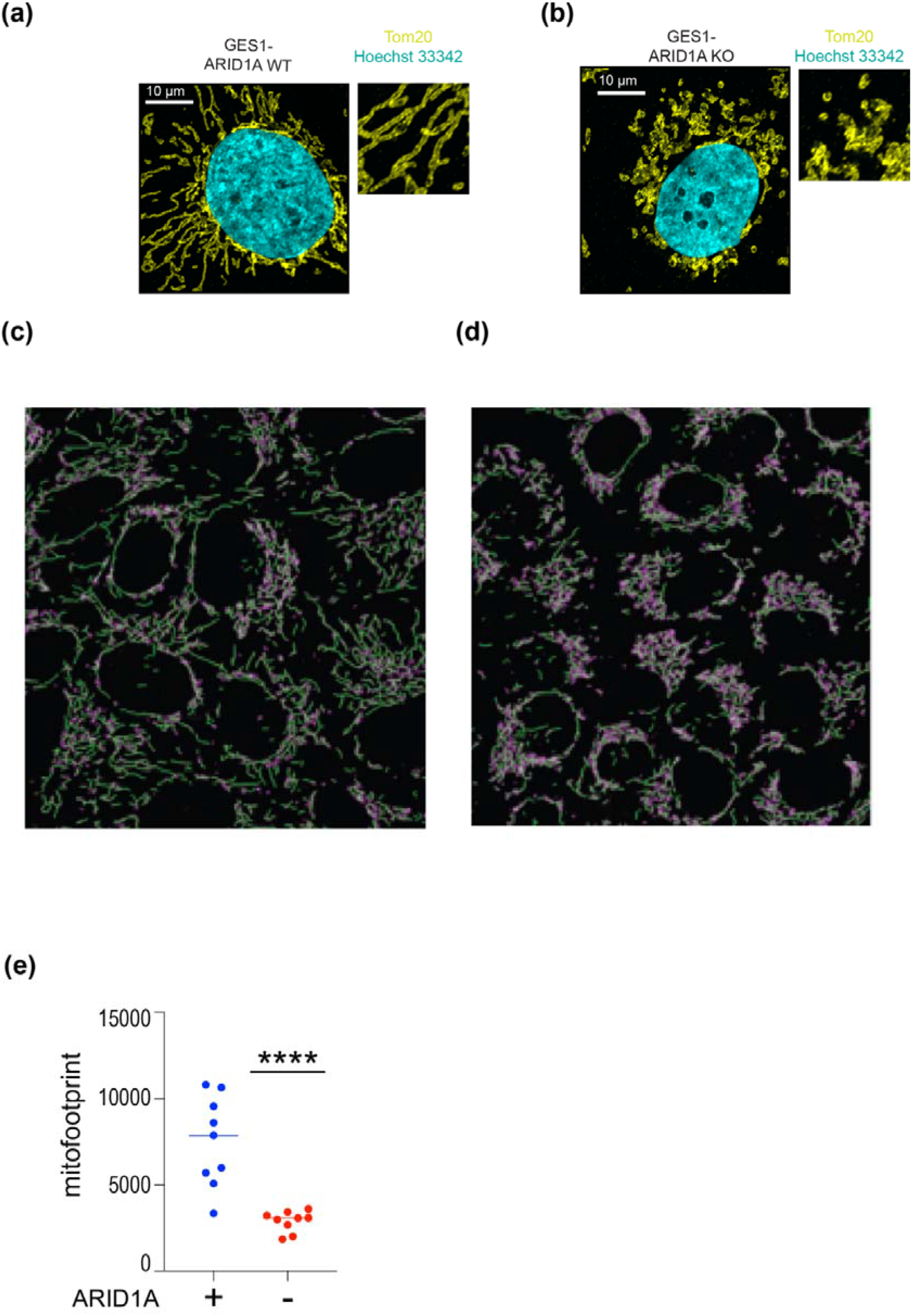
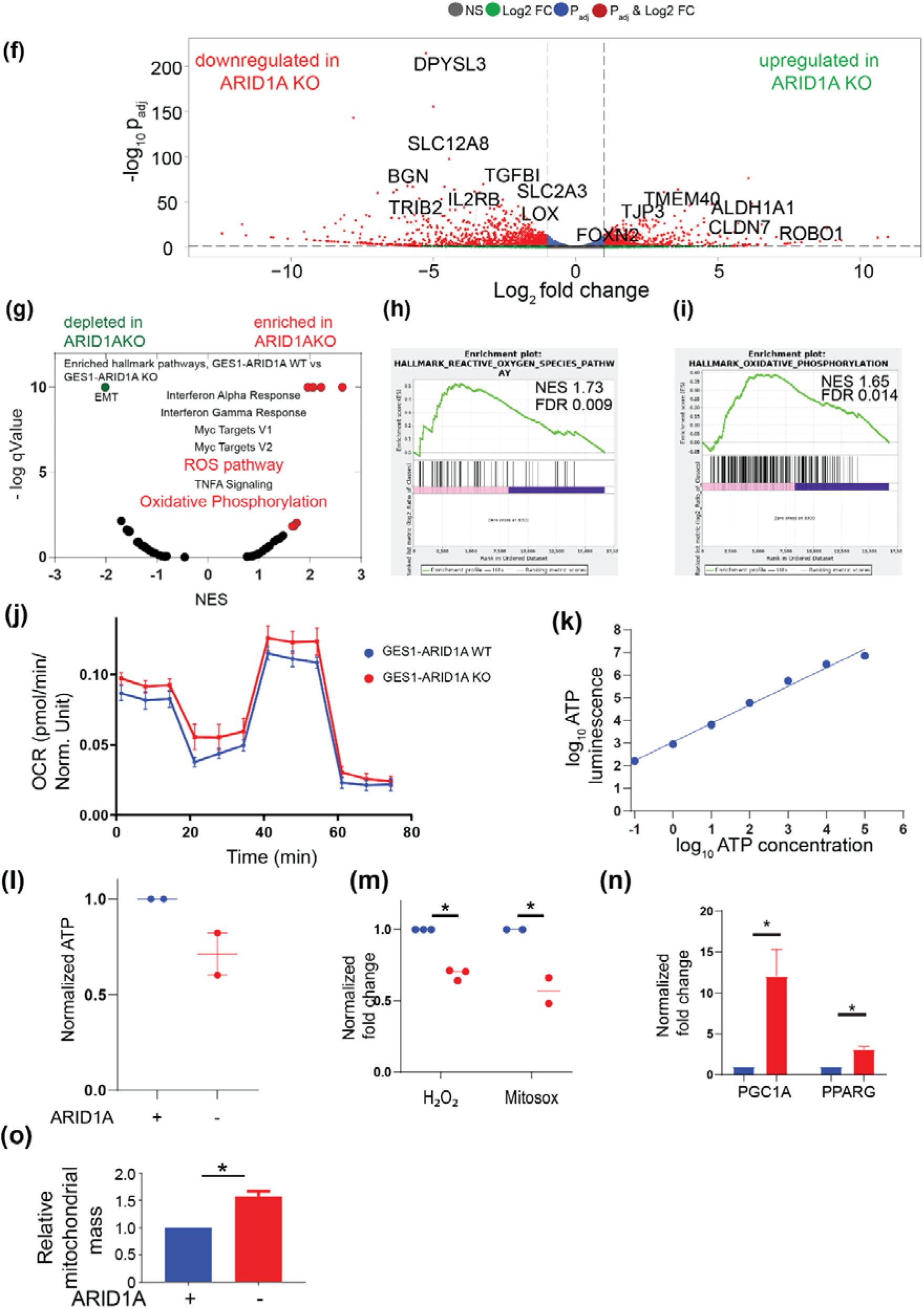
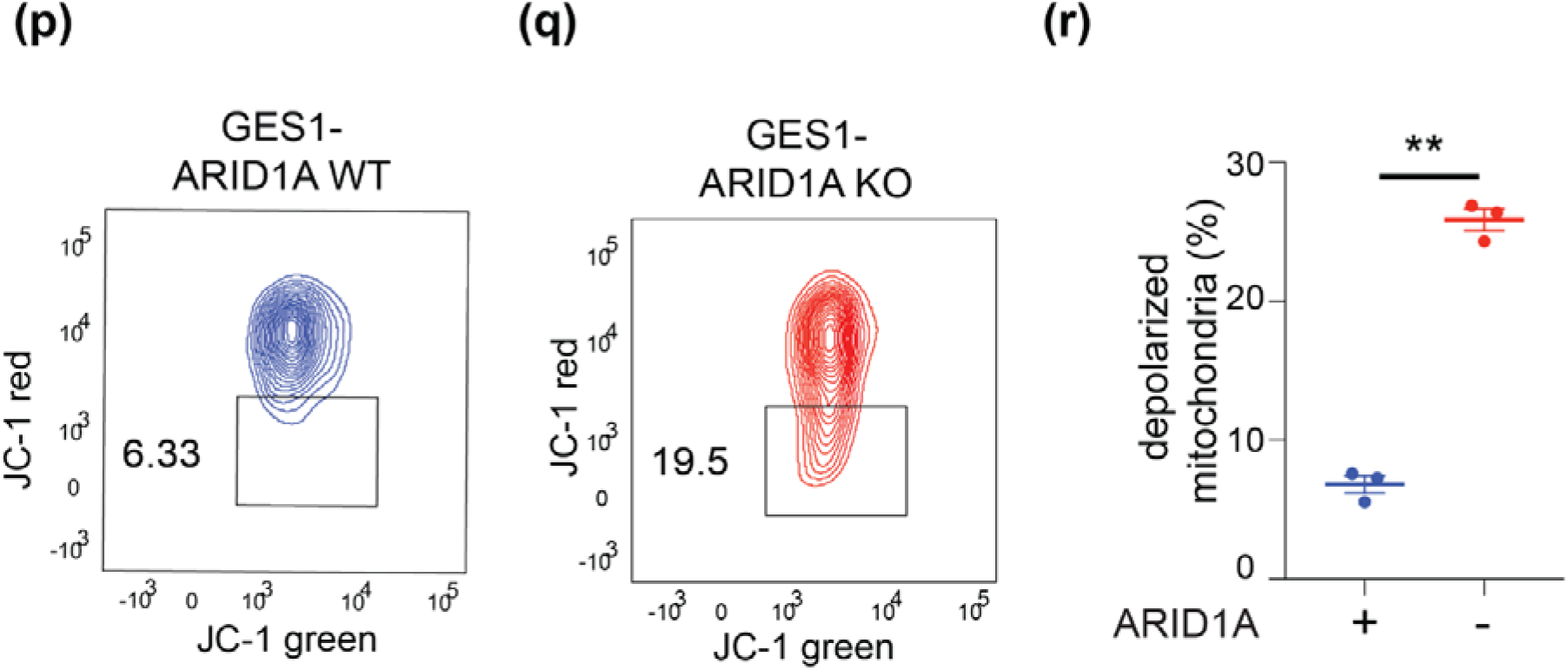
ARID1A KO cells have a grossly abnormal mitochondrial phenotype. Confocal images (63x, Airyscan) of GES1-ARID1A WT and GES1-ARID1A KO cells stained with mitochondrial marker Tom20 (yellow) and counterstained with Hoechst33342 (cyan) showing different mitochondrial morphology between - a) ARID1A-WT (filamentous) and b) ARID1A-KO (globular). The images in the inset are at 5x digital zoom. Confocal images of GES1-ARID1A WT and KO cells stained with Tom20, showing difference in mitochondrial morphology; ARID1A-WT (left) – filamentous mitochondria, ARID1A-KO (right) – globular. The images on the right represent segmentation obtained from MiNA algorithm (Image J) used for quantitating mitochondrial phenotype. c) GES1-ARID1A WT. d) GES1-ARID1A KO. e) Comparison of the mitochondrial footprint between GES1-ARID1A WT and GES1-ARID1A KO cells demonstrating the KO cells have a lower mitochondrial footprint. f) Volcano plot showing differentially expressed genes from RNAseq analysis (n=2 replicates) of GES1-ARID1A WT and KO cells, g) GSEA hallmark pathway enrichment analysis between GES1-ARID1A WT and KO cells showing enrichment in ROS and Oxidative phosphorylation pathways, h) GSEA enrichment plot with normalized enrichment score (NES) for ROS pathway between GES1-ARID1A WT and KO cells. i) GSEA enrichment plot with normalized enrichment score (NES) for oxidative phosphorylation pathway between GES1-ARID1A WT and KO cells. j) Sea horse experiment showing increased basal respiration in GES1-ARID1A KO cells as well as increased maximal respiratory capacity compared to GES1-ARID1A WT cells. k) Standard curve for ATP determination used in l) demonstrating the validity of the assay: The amount of ATP strongly correlates with the luminescence produced. l) Dot plot showing no significant change in ATP levels between GES1-ARID1A WT and GES-1-ARID1A KO cells m) Comparison of total H_2_O_2_ levels and mitochondrial ROS measured in GES1-ARID1A WT (blue) and KO (red) cells. n) qPCR analysis of GES1-ARID1A WT (blue) and GES1-ARID1A KO (red) cells for PGC1A, a regulator of mitochondrial biogenesis and PPARG showing increased expression in GES1-ARID1A KO cells. Student’s t-test, p<0.05. o) qPCR analysis of GES1-ARID1A WT and GES1-ARID1A KO cells showing increased total mitochondrial mass estimated by qPCR based expression of nuclear and mitochondrial encoded genes. The genes used were as following – mitochondria encoded cytochrome B, mitochondria encoded CoxI, mitochondria encoded ATP6, nuclear encoded ATP5B, nuclear gene desert region chromosome 10. p) Representative flow cytometry plots of JC-1 staining measuring mitochondrial membrane potential in GES1-ARID1A WT cells. The reduction in red fluorescence from JC-1 is indicative of cells with reduced mitochondrial membrane potential. q) Representative flow cytometry plots of JC-1 staining measuring mitochondrial membrane potential in GES1-ARID1A KO cells. r) Quantitation of depolarized mitochondria in JC-1 flow cytometry experiments (n=3). Error bars represent SEM of 3 independent biological experiments, p<0.01, student’s t-test.

ARID1A is a nuclear protein that functions as a chromatin-remodeling complex to influence transcription. We performed an unbiased whole transcriptomic analysis comparing GES1-ARID1A WT and KO cells to look for mitochondrial pathways affected by changes in the transcriptomic landscape upon ARID1A loss. Unsurprisingly, the loss of ARID1A dramatically rewired the transcriptomic landscape, resulting in a substantial downregulation of target genes (Fig. 4F). This is in line with the localization of ARID1A to actively transcribed genomic regions(14). While gene enrichment analysis revealed increases in interferon genes and genes modulated by MYC in ARID1A KO cells (Fig. 4G), we were particularly interested in an upregulation of oxidative phosphorylation and reactive oxygen species (ROS) pathways (Fig. 4H and 4I). In light of mitochondrial phenotype and enriched ROS and OXPHOS pathways in ARID1A KO cells, we next investigated the oxygen consumption rate in these isogenic cell lines. We found that ARID1A KO cells indeed had a higher basal oxygen consumption (Fig. 4J), in keeping with a prior report that loss of ARID1A leads to an increased dependence on mitochondrial oxidative phosphorylation in cancer cells(24). Interestingly, all of the 13 proteins encoded by the mitochondrial genome (and translated by the mitochondrial ribosome) are considered important for oxidative phosphorylation(24,25). As the results obtained from our CRISPR screen revealed that sensitivity to PLK1 inhibition in ARID1A KO cells was dependent on the expression of mitochondrial translational machinery (Fig. 3C, 3D and 3E), we hypothesized that the increased oxidative phosphorylation in ARID1A deficient cells leads to increased reliance on PLK1 function, in a manner dependent on mitochondrial protein translation.

Interestingly however, when measuring ATP production in these cells, we noted that the increased oxygen consumption was not accompanied by increased ATP production (Fig. 4K and 4L). Next, we measured total H_2_O_2_ and mitochondrial ROS (Fig. 4M) and found ARID1A KO cells actually accumulated lower levels of these ROS moieties(26), in line with an uncoupling of oxygen consumption from its subsequent metabolism. This increased oxygen consumption is accompanied by heightened mitochondrial biogenesis, as evidenced by increased expression of the biogenesis proteins PGC1A and PPARG(27) (Fig. 4N), as well as increased mitochondrial mass in ARID1A KO cells (Fig. 4O). The enrichment of ROS and oxidative phosphorylation pathways noted on RNAseq is consistent with findings that cells compensate for increased energy needs by increasing mitochondrial mass(28). The increase in mitochondrial biogenesis and oxygen consumption without commensurate increases in ATP production in ARID1A deficient cells suggests that these are compensatory mechanisms to an underlying metabolic defect in their mitochondria. When examining the mitochondrial membrane potential(29) of these cells using the JC-1 dye, we note that a significant proportion of ARID1A KO cells have lower mitochondrial membrane potential compared to the WT cells, confirming a baseline loss of mitochondrial fitness (Fig. 4P, 4Q, 4R).

### PLK1 localizes to mitochondria in interphase cells

Given the loss of mitochondrial fitness in ARID1A deficient cells, and the reliance of ARID1A KO cells on PLK1 function, we then asked whether PLK1 localized to mitochondria in interphase cells. We validated an antibody for PLK1 immunofluorescence using siRNA knockdown of PLK1, and demonstrated that the dispersed cytoplasmic signal was indeed PLK1 (Fig. 5A). Using confocal microscopy, we demonstrated that several of these signals in interphase cells co-localized with the mitochondrial marker Tom20 (Fig. 5B). We then explored PLK1 localization to mitochondria using expansion microscopy, a super-resolution like technique. This technique utilizes a polymer to symmetrically enlarge biological samples by up to 10-fold, thereby allowing clear visualization of protein localization to sub-micron subcellular structures(30). Using this method, we noted that signals of PLK1 did localize in structures staining for ATP5B and ATP5A, subunits of complex V of the mitochondrial electron transport chain (Fig. 5C and 5D). We then cross-validated these findings by biochemical fractionation of mitochondria from cellular extracts, showing that PLK1 was present in the mitochondrial fraction (Fig. 5E). PLK1 does not have a canonical mitochondrial localization signal, and these experiments do not distinguish if the protein is within or merely tethered to the mitochondria. However, taken together they do provide evidence that a fraction of cytoplasmic PLK1 is associated with mitochondria, and could directly play a role in regulating mitochondrial function.

**Fig. 5:**
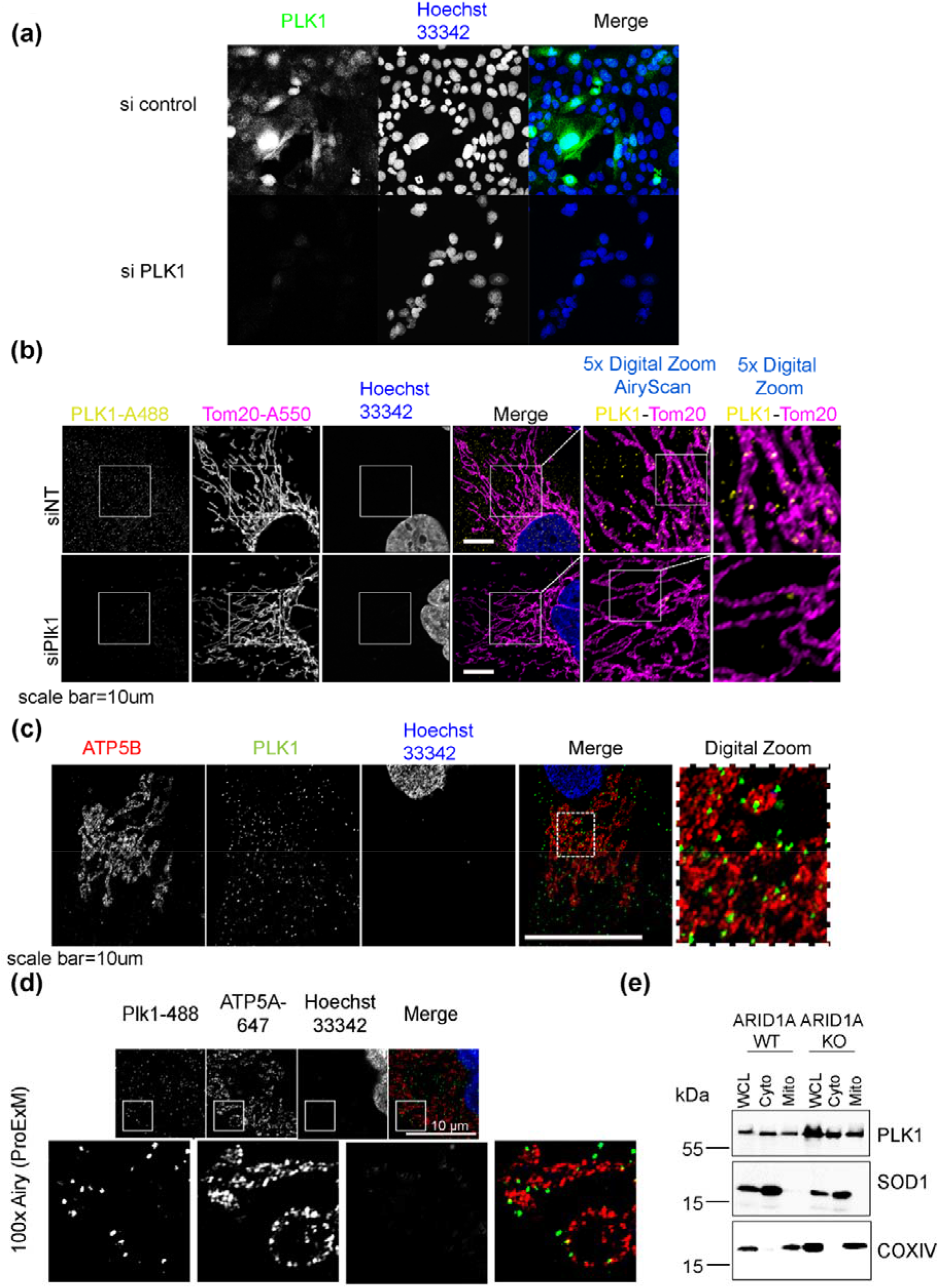
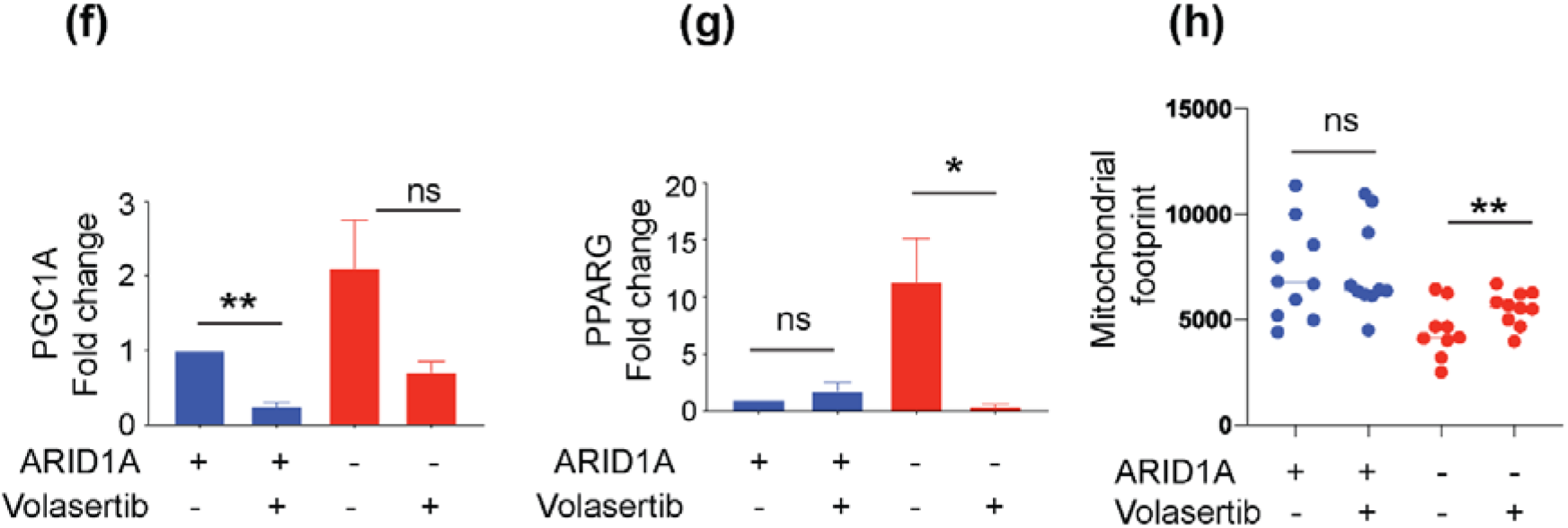
PLK1 localises to the mitochondria. a) Validation of PLK1 antibody specificity. Confocal images showing PLK1 staining in GES1-ARID1A WT cells transfected with sicontrol and siPLK1. b) Confocal images of GES1-ARID1A WT cells with and without siPLK1 showing PLK1 localization to mitochondria. Tom20 is used as a mitochondrial marker, and DNA is stained using Hoechst 33342. siPLK1 is used as a quality control for antibody specificity. c) Confocal images of GES1-ARID1A WT cells showing PLK1 localization to mitochondria. ATP5B is used as a mitochondrial marker, and DNA is stained using Hoechst 33342. d) Expansion super-resolution microscopy images of ATP5A and PLK1 showing localization of PLK1 in mitochondria. e) Biochemical fractionation of mitochondria showing PLK1 localization in mitochondria. COX IV was used as a control for mitochondrial fraction while SOD 1 was used for cytoplasmic fraction. f) qPCR analysis of PGC1A in Volasertib treated/untreated, GES1-ARID1A WT and GES1-ARID1A KO cells. Volasertib treatment decreases the expression of PGC1A in GES1-ARID1A KO cells. g) qPCR analysis of PPARG in Volasertib treated/untreated, GES1-ARID1A WT and GES1-ARID1A KO cells. Volasertib treatment decreases the expression of PPARG in GES1-ARID1A KO cells. h) Mitochondrial footprint (volume) analysis for GES1-ARID1A WT and GES1-ARID1A KO cells. Volasertib treatment does not increase mitochondrial footprint in the WT cells but increases it in the KO cells.

As we noted ARID1A deficient cells to have higher mitochondrial biogenesis, we hypothesized that PLK1 may be critical to maintain this state. We therefore evaluated whether the high expression of the mitochondrial biogenesis regulators PGC1A and PPARG were dependent on PLK1. Indeed, addition of Volasertib dramatically reduced the expression of both genes in the ARID1A KO cells (Fig. 5F and 5G). We next asked if Volasertib treatment effected mitochondrial morphology as measured by the mitochondrial marker Tom20. Treatment with Volasertib did not significantly change the mitofootprint in ARID1A WT. However, it did result in a small but statistically significant increase in the mitofootprint in the ARID1A KO cells (Fig. 5H). Together these results suggest that the aberrant mitochondrial biogenesis phenotype in ARID1A deficient cells is dependent (at least partially) on PLK1 function.

### PLK1 inhibition exacerbates the mitochondrial fitness defect in ARID1A deficient cells

RNAseq analyses of ARID1A WT and KO cells in the presence of Volasertib revealed a similar gene set differences as without Volasertib (Fig. S2A), although the enrichment scores for both oxidative phosphorylation and ROS pathways were now higher after Volasertib addition (Fig. S2B and S2C). We therefore tested if the alteration in mitochondrial biogenesis induced by PLK1 inhibition could impact on overall mitochondrial metabolism and respiration in the ARID1A KO cells. Interestingly, 1D NMR revealed marked differences in the metabolome of GES1-ARID1A KO cells when compared to WT cells, with this difference accentuated by the addition of Volasertib. (Fig. 6A, and Fig. S3). Measurements of oxygen consumption using the seahorse assay revealed that ARID1A KO cells have significantly higher basal respiration (Fig. 6B), in line with earlier observations (Fig. 4J)(24). Treatment with Volasertib however dramatically increased oxidative phosphorylation in ARID1A KO cells compared to WT cells. These results suggest that PLK1 mediated mitochondrial biogenesis is important to maintain the “altered normal” state of metabolism and respiration in ARID1A KO cells. We again investigated ATP levels in the isogenic cell lines as before, but this time in the presence of Volasertib, and found no significant effect of the treatment on the ATP levels between the isogenic cell lines (Fig. 6C). We therefore hypothesized that the dramatically increased demands on metabolism and in ARID1A deficient cells in the setting of PLK1 inhibition would have an effect on the membrane potential of mitochondria, which were already compromised in ARID1A KO cells. PLK1 inhibition by Volasertib significantly increased the fraction of depolarized mitochondria in ARID1A deficient cells, while only a mild increase was seen in ARID1A WT cells (Fig. 6D and Fig. 6E). This was accompanied by an increase in the apoptotic markers cleaved caspase 3 and cleaved PARP (Fig. 6F).

**Fig. 6:**
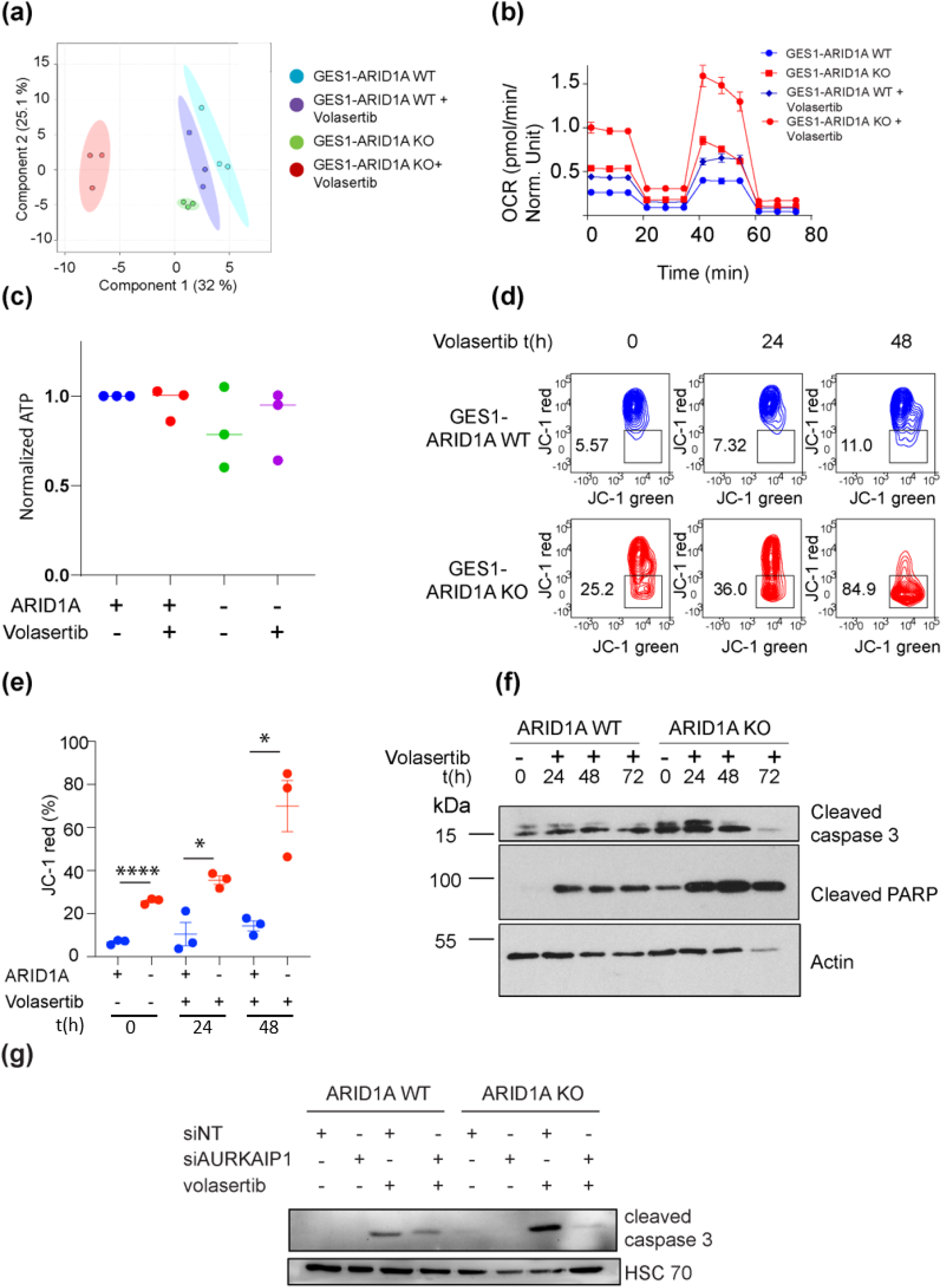
PLK1 inhibition differentially affects the mitochondria of ARID1A KO cells. a) PCA plots from NMR-based unbiased metabolomics of GES1-ARID1A WT and GES1-ARID1A KO cells, with and without Volasertib. There is a drastic change in the metabolome of ARID1A KO cells upon treatment with Volasertib. b) Sea horse experiment showing increased basal respiration in GES1-ARID1A KO cells as well as increased maximal respiratory capacity compared to GES1-ARID1A WT cells. Addition of Volasertib further exacerbates this phenotype in the GES1-ARID1A KO cells. c) Dot plot showing no significant change in ATP levels between GES1-ARID1A WT and GES-1-ARID1A KO cells. Quantification representing 3 independent biological experiments (n=3). d) Representative images of JC-1 assay in GES1-ARID1A WT and GES1-ARID1A KO cells treated with Volasertib for 24 and 48 h respectively. Measurements were obtained in BD FACS. e) Quantitation of percentage of cells showing more depolarization of mitochondria in ARID1A KO cells (lower mitochondrial membrane potential). Treatment with Volasertib increased depolarization with time and to a greater extent in GES1-ARID1A KO cells. Error bars represent SEM of 3 independent biological experiments, **** = p<0.0001, * = p<0.05, student’s t-test. f) Western blot analysis of GES1-ARID1A WT and GES1-ARID1A KO cells with Volasertib treatment for 24, 48 h and 72 h. GES1-ARID1A KO cells show higher apoptosis at the baseline and post Volasertib treatment indicated by higher levels of cleaved caspase 3 and cleaved PARP. Actin was used as loading control. g) Western blot analysis of GES1-ARID1A WT and GES1-ARID1A KO cells transfected with siNT/ siAURKAIP1. AURKAIP1 is a mitochondrial translation gene (see Fig 3e). Knockdown of AURKAIP1 reduces Volasertib-induced apoptosis in ARID1A KO cells.

To directly prove that mitochondrial function was critical for Volasertib induced apoptosis in ARID1A deficient cells, we knocked down AURKAIP1 (the top-hit in the mitochondrial translation pathway in our CRISPR screen), in the setting of volasertib treatment. siAURKAIP1 was able to rescue ARID1A KO cells from Volasertib induced apoptosis (Fig. 6G). Taken together, these data suggest that ARID1A deficient cells have a metabolic defect characterized in increased oxygen consumption but without the commensurate increase in ATP production. This phenomenon is exacerbated by PLK1 inhibition (Fig. 7A). The survival of cells with this metabolic defect is dependent on non-canonical PLK1 activity that is ostensibly important for maintaining processes that rely on or involve the mitochondrial ribosomal translation machinery.

**Fig. 7:**
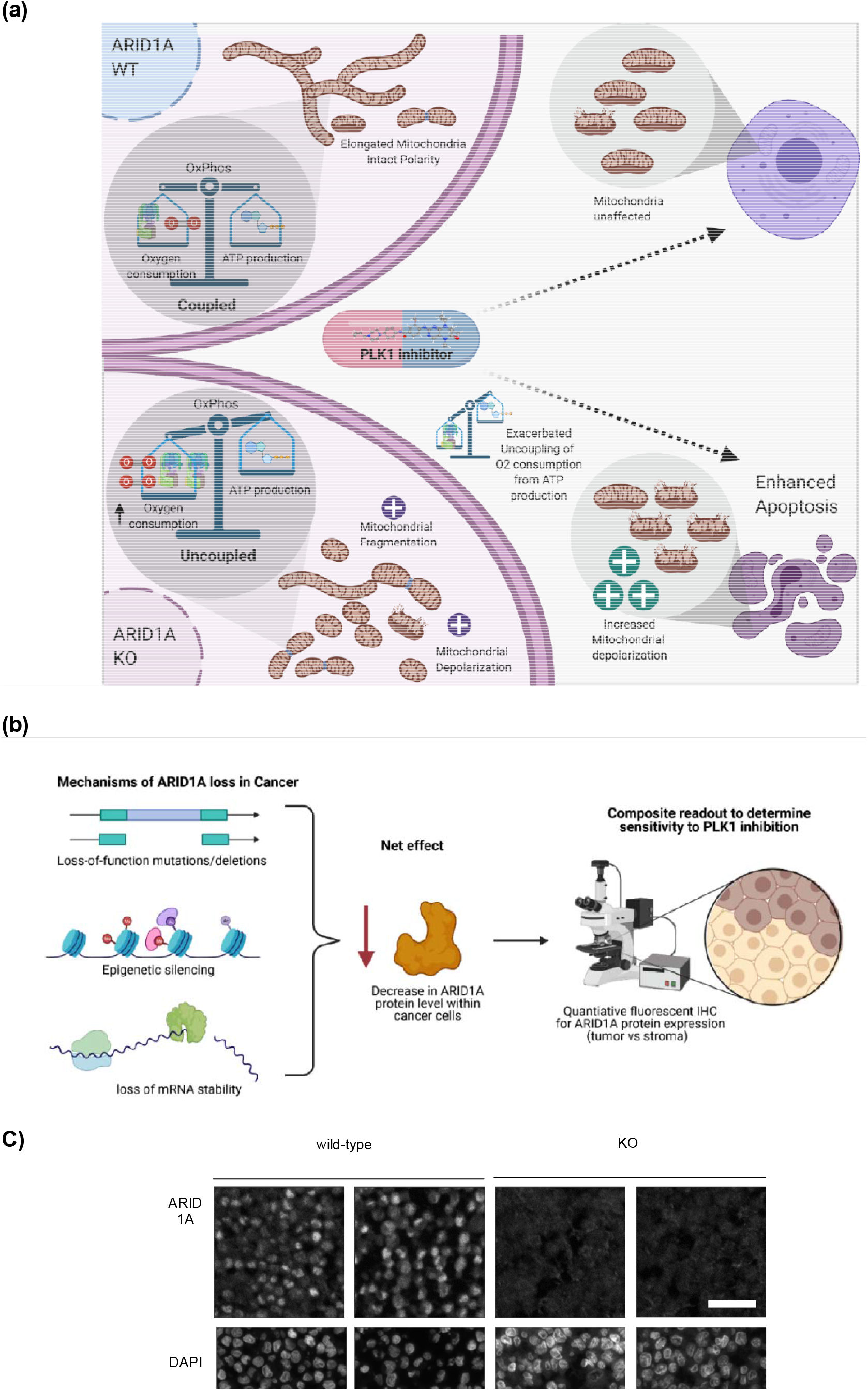
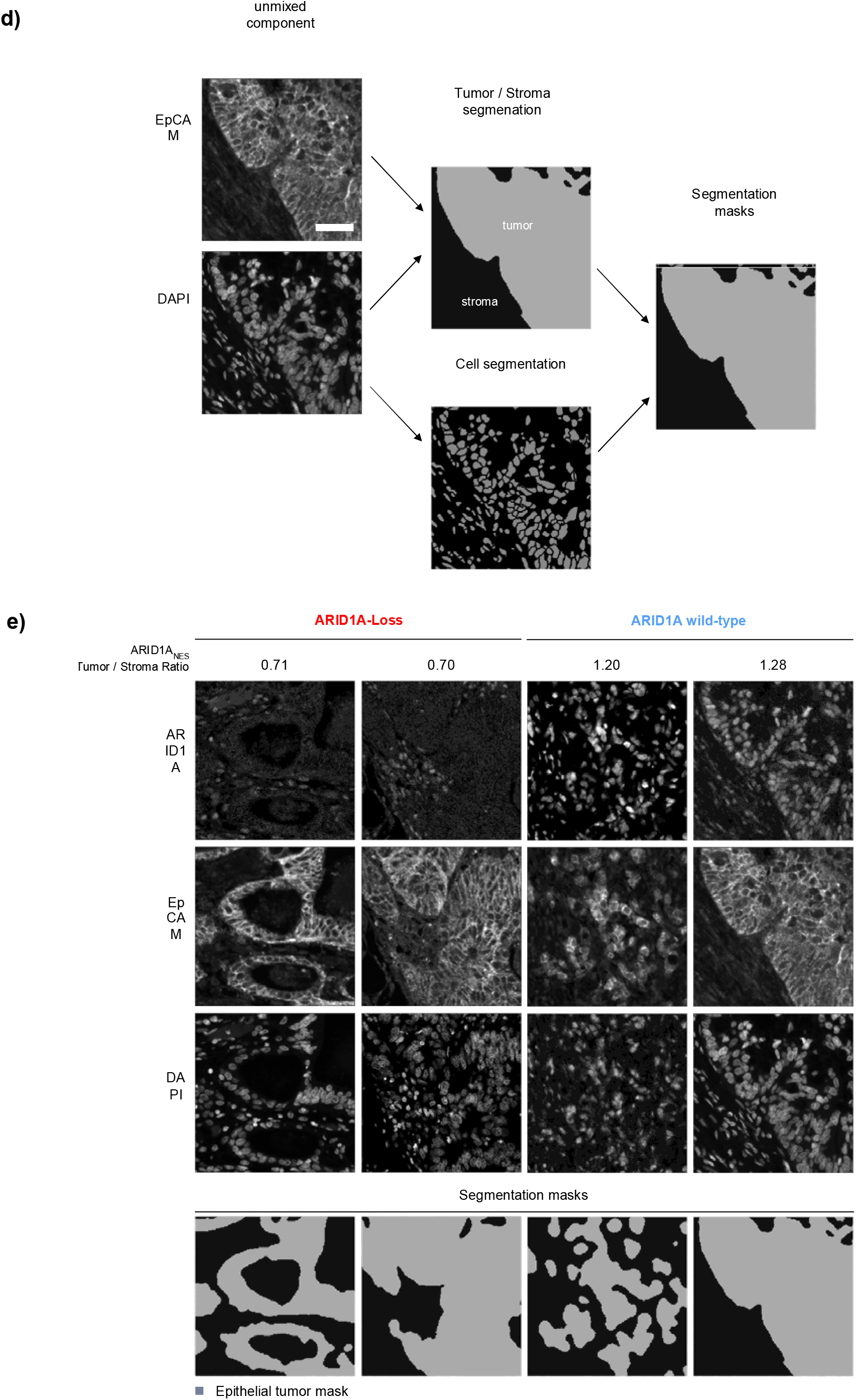
Schematic representation of the uncoupling of oxidative phosphorylation and ATP production in ARID1A KO cells and the subsequent consequences of PLK1 inhibition. a) Cartoon illustrating the mechanism for differential sensitivity of ARID1A deficient cells to PLK1 inhibition. ARID1A deficient cells display an uncoupling of oxygen consumption from ATP production at the baseline state, and this is worsened when PLK1 is inhibited-leading to increased mitochondrial depolarization and apoptosis. b) Overview of the principle underlying an ARID1A quantitative IHC assay to predict PLK1i sensitivity. c) Validation of an anti-ARID1A antibody specificity for formalin-fixed samples; using OVCAR-3 ARID1A-expressing and knock-out (KO) formalin-fixed cell blocks. d) Schematic representation of machine-learning assisted analysis of ARID1A multiplexed fluorescent IHC samples, using EpCAM as an epithelial marker to differentiate epithelial tumor areas from stroma. e) Examples of unmixed monochrome images of gastric cancer samples with ARID1A-Loss (left) and intact ARID1A (right). ARID1A is retained within the stroma of ARID1A deficient cancers, and serves as an internal control. Quantitative ARID1A_NES_ Ratio between Tumor / Stromal compartments is indicated above the images. Cell segmentation and tissue segmentaion masks were created based on unmixed EpCAM and DAPI nuclear counterstaining images. DAPI - 4′,6-diamidino-2-phenylindole. Scale bar is 50μm in all panels.

### Establishment of a quantitative IHC assay to select for PLK1 inhibition therapy

Our results establish ARID1A loss in cancer as a possible biomarker for sensitivity to PLK1 inhibition. To translate this finding into the clinic, it is imperative that a robust ARID1A detection assay be developed (Fig. 7B). As there are multiple ways in which ARID1A loss can occur (including loss of function mutations, epigenetic suppression and aberrant mRNA processing), the measurement of ARID1A at the protein level is the most comprehensive way to assess ARID1A loss (Fig. 7B). In order to develop a quantitative assay of ARID1A protein loss, we first validated a rabbit monoclonal antibody (ab182561) for reliable detection of ARID1A expression in FFPE samples. The antibody recognized a clear nuclear signal in FFPE cell-blocks (consistent with ARID1A function as chromatin remodeler), which was abrogated in knock-out cell blocks (Fig. 7C). We have previously used a “nuclear expression score” (31) as an overall assessment of a given protein per case, which is reflective of the average intensity of a given protein across all imaged cells. We similarly established an ARID1A nuclear expression score (ARID1A_NES_) as a starting point for this assay. As ARID1A loss is frequently observed in tumor cells, but not stromal cells(32) we used a marker of epithelial cells to differentiate the quantitative analysis between tumor and stromal cells (Fig. 7D). We hence used a Tumor / Stroma ARID1A_NES_ Ratio as a readout for ARID1A loss (Fig. 7E). In our proof-of-concept assessment, gastric cancer cases with a visually appreciable loss of ARID1A protein expression, show a Tumor / Stroma ARID1A_NES_ Ratio < 1 (Fig. 7E). We propose that this method could be widely applied across cancer types for the identification of cases with loss of ARID1A expression in tumor cells, and therefore likelihood of response to PLK1 inhibition.

## Discussion

In this study, we show that PLK1 inhibitors are preferentially lethal to cells with loss of the tumor suppressor ARID1A, through a cell-cycle independent mechanism. Although the canonical functions of PLK1 promote G2/M cell cycle transition, there is increasing data on its role in modulating metabolic states (33–35), and in maintaining other cells with dysfunctional mitochondria (36). This appears to also be true in the setting of ARID1A loss, where the cells have markedly aberrant mitochondrial biogenesis with a phenotype of extensive fragmentation, and are highly sensitive to PLK1 inhibition. The altered biogenesis in ARID1A deficient cells is likely a consequence of altered metabolism, as mitochondrial form and function are intricately linked (37,38). It is known that ARID1A and SWI/SNF loss alters cellular metabolism causing an increased reliance on oxidative phosphorylation (OXPHOS)(24,39). We now describe that ATP production is uncoupled from increased baseline oxygen consumption and expression of OXPHOS genes in the setting of ARID1A loss; i.e. despite showing increased oxygen consumption rate (OCR), ARID1A deficient cells produce less ATP than their isogenic counterparts. This phenotype is not lethal, as ARID1A deficient cells are able to proliferate, albeit with the abovementioned abnormal morphology, suggesting that the increases in oxygen consumption and OXPHOS gene expression are compensatory mechanisms to allow survival of these cells. Interestingly, upon chemical inhibition of PLK1 in ARID1A deficient cells, <u>oxygen consumption is further “uncoupled” from ATP production</u>, with markedly increased OCR but without additional ATP production. This is accompanied by extensive mitochondrial depolarization/ apoptosis. Conversely, in normal cells, PLK1 inhibition has a minimal effect on oxygen consumption, and leads to cell death only at higher doses through canonical G2/M arrest. These results suggest that ARID1A deficient cells are non-epistatically dependent on a PLK1-mediated mechanism that regulates oxygen consumption rate.

We also show for the first time that a fraction of cellular PLK1 localizes to mitochondria, adding weight to the evidence for its direct role at the organelle. Aurora kinase A, which directly phosphorylates PLK1, was previously shown to localize to mitochondria and regulate mitochondrial biogenesis(40). As proteins with key roles in orchestrating the fidelity of normal cell division(41), it is possible that the Aurora kinase A – PLK1 axis is active in mitochondria and plays a role in coordinating mitochondrial function and biogenesis with cell division. Our work lays out the rationale for additional studies to map specific PLK1 substrates in the mitochondria, which may help understand how the coordination of metabolism, mitochondrial division and cell division is maintained.

Our current work however highlights increased reliance on mitochondrial protein synthesis as an Achilles heel in ARID1A deficient cells; rendering them sensitive to PLK1 inhibition. Our CRISPR screen highlighted mitochondrial ribosomal proteins as critical for Volasertib induced death in ARID1A deficient cells. Mitochondrial ribosomal proteins are essential for the synthesis of OXPHOS components, and their depletion or malfunction is usually associated with reduced oxygen consumption (42). Together with the finding of increased oxygen consumption after PLK1 inhibition in ARID1A deficient cells, we propose a model (Fig. 7A) where PLK1 limits oxygen consumption in ARIDA1 deficient cells through controlling one or more mitochondrially translated proteins. These findings are concordant with retrospective analysis of a phase 1 trial of the PLK1 inhibitor, Onvansertib, which revealed an association between tumor shrinkage and increased expression of OXPHOS genes(43) (as is noted in ARID1A deficient tumors and cells).

Of clinical importance, our work outlines a clear translational path in cancer for the development of PLK1 inhibitors. ARID1A is frequently mutated gene in a wide variety of epithelial and haematological cancers. These mutations leading to loss of protein(44) are even more frequent in certain subtypes such as gastric cancer (~20%) and ovarian clear-cell carcinoma (~40%)(45,46). ARID1A deficient gastric cancers have worse clinical outcomes than ARID1A proficient cancers(17). In view of its high mutational frequency, ARID1A loss has emerged as an attractive target for exploring synthetic lethal interactions. While several candidates have emerged from pre-clinical studies(8,13,18,21,47,48), to date none of these are established as clinical strategies to treat patients with ARID1A mutant cancers. In our knockout cell models, Volasertib was more potent at killing ARID1A deficient cells than Olaparib, ATR inhibitors or Dasatinib (Fig S2), further highlighting its utility in this setting.

Volasertib has not had widespread clinical success despite its initial promising activity, predominantly due to its toxicity and the lack of a biomarker for selection of cases. While the dose limiting haematological toxicity of these inhibitors is likely due to their effect on cell-cycle progression/ stem-cell proliferation at high doses, we offer rationale for a therapeutic window to use lower doses that induce mitochondrial depolarization and apoptosis in ARID1A deficient tumor cells, without significant effects on the cell cycle of non-tumor cells. This could form the scientific basis for the testing of Volasertib or other PLK1 inhibitors in a basket trial of cancers with ARID1A loss, potentially at doses that are well tolerated clinically. The quantitative assay that we describe to measure ARID1A expression at the protein level would form a broad selection marker for the inclusion of cases with a wide range of genetic and epigenetic mechanisms leading to loss of ARID1A.

## Author contributions

Conceptualization: USS, MMH, ADJ

Data curation: USS, OA, AS, HY

Formal analysis: USS, SL, MMH

Funding acquisition: ADJ

Investigation: USS, GKR, JDW, MYL, PCP, BWQT, LH, RJ, KS, MB, SC, TSCN, YP, SL

Methodology: USS, DK, SC, WLT, CF, ADJ

Project administration: ADJ

Resources: LHKL, SP, KC, PT, DK, YWP, CF, DSPT, WLT, ADJ

Software – OA, HY

Supervision: ADJ

Validation: USS, LHKL, SP, KC, PT, DK, YWP, CF, DSPT, WLT, AS, HY, OA

Visualization: USS, MMH, ADJ

Writing – original draft – USS, ADJ

Writing – review and editing - LHKL, SP, KC, PT, DK, YWP, CF, DSPT, WLT, PJ, ADJ

## Notes

**Financial Support:** ADJ is supported by the Singapore Ministry of Health’s National Medical Research Council Transition Award (NMRC/TA/0052/2016). Work in ADJ’s laboratory is also funded by the Cancer Science Institute of Singapore, through the National Research Foundation Singapore and the Singapore Ministry of Education under its Research Centres of Excellence initiative. Part of this work was funded through a collaborative grant between ADJ and DSPT from the Singapore Ministry of Health’s National Medical Research Council (NMRC/CIRG/1400/2014).

**Conflicts of Interest:** ADJ; honoraria from AstraZeneca, Janssen and MSD, travel funding from Perkin Elmer, and research funding from AstraZeneca and Janssen. DSPT; honoraria from AstraZeneca, Roche, Bayer, MSD, Merck Serono, Tessa Therapeutics, Novartis, and Genmab and research funding from AstraZeneca, Bayer and Karyopharm. The other co-authors have no conflicts of interest to declare.

### Competing Interest Statement

ADJ; honoraria from AstraZeneca, Janssen and MSD, travel funding from Perkin Elmer, and research funding from AstraZeneca and Janssen. DSPT; honoraria from AstraZeneca, Roche, Bayer, MSD, Merck Serono, Tessa Therapeutics, Novartis, and Genmab and research funding from AstraZeneca, Bayer and Karyopharm. The other co-authors have no conflicts of interest to declare.

