## Supplementary Methods, Tables and Figures for "PLK1 inhibition selectively kills ARID1A deficient cells through uncoupling of oxygen consumption from ATP production"

**Supplementary Materials**

**Supplementary Methods**

### Colony formation and cell proliferation assays (crystal violet)

For colony formation assay, 200 cells per well were plated in a 6 well plate, and treated with the indicated concentration of chemotherapeutics for 14 days. Cells were washed with 3xPBS and stained with crystal violet followed by washing with water to remove excess stain. Images were obtained under microscope using axiovision software. Colonies formed were manually counted and plotted in PRISM^TM^. For cellular proliferation, 800 cells per well were plated in 6 well plates and treated with indicated concentrations of chemotherapeutics every alternate day for 10 days. The cells were stained with crystal violet and images taken as described above.

Xenograft mice experiments

OVCAR3 wild type cells (5 × 10^6^ cells) or OVCAR3 ARID1A knock out cells (5 × 10^6^ cells) were injected subcutaneously in the right flank of female NOD/MrkBomTac-Prkdcscid mice  (InVivos, Singapore.). When the xenografts had reached an average volume of 300 mm^3^ animals were randomized into four groups of five mice each. Volasertib was formulated in 90% corn oil and injected via intraperitoneal route. The dosage of Volasertib administered was 15 mg/kg of body weight. The results were converted to tumor volume (mm^3^) by the formula π(length × width^2^)/6. The weight of the mice was determined using an electronic balance as an indicator of drug toxicity. The animal experiments were conducted at National University of Singapore in accordance with the Institutional Guidelines for the Care and Use of Laboratory Animals.

### RNA-Seq analysis and gene expression profiling

Gene expression profiling, pathway enrichment analysis and GSEA from RNA-Seq data have been performed by using CSI NGS Portal(1) (<https://csibioinfo.nus.edu.sg/csingsportal>). Briefly, raw fastq files were trimmed by using Trimmomatic(2) for adapter removal. The clean reads were aligned to the reference human genome (hg19) by using STAR(3) (v2.7.3a) with default parameters. The gene expression quantification was done by using HTSeq-count(4) (v0.11.2) in strand-specific mode “-s reverse” to obtain raw read counts for each gene, and read counts only from the sense strand are used. First, PCA and hierarchical clustering were performed with regionReport(5) to compare the gene expression profiles of the samples, by using the top 500 genes with the highest variance across the samples. Then the differential gene expression analysis was performed by using DESeq2(6) (v1.24.0) starting from the raw read counts by comparing the samples from GES1-ARID1A KO to GES1-ARID1A WT after collapsing the three samples within each group as replicates. The genes that are not expressed or lowly expressed (read counts <= 2 on average per sample) were removed from the analysis. In total, 258 and 715 genes were significantly up- and down-regulated, respectively (log_2_fold change>2, p_adj_value<0.05), in the KO samples compared to the WT samples, after correction for multiple hypothesis testing (P_adj_ < 0.05, using Benjamini-Hochberg method). These differentially expressed genes were used for the pathway enrichment analysis (ReactomePA database(7)). Additionally, gene set enrichment analysis (GSEA(8)) was performed by using the gene expression data for all the genes normalized by DESeq2(6). To be comprehensive, all pathways and gene sets were selected as the input for the enrichment analyses. The significantly altered pathways were identified and plotted as barplot by using R packages ReactomePA(7) and enrichplot(9), respectively.

### 1D NMR Metabolomics:

Cells were plated in 2x10 cm dishes and incubated overnight at 37°C in 5% CO_2_ followed by treatment with DMSO/10nM Volasertib for 24 hours. Post treatment cells were washed 2x with ice cold PBS and fixed with 100% ice cold methanol and harvested. Cells were then lysed using a probe sonicator on ice. The lysate was kept at -20°C for 30 min to promote protein precipitation and centrifuged at 15,000 rpm for 30 min at 4°C. The supernatant was collected and dried under vacuum at 30^o^C. The dried metabolite samples were resuspended in PBS prepared in heavy water (D_2_O) containing DSS (internal standard) followed by NMR analysis.

One-dimensional ^1^H NMR spectrometry (DRX500, Bruker USA) was performed and NMR spectral data was Fourier-transformed followed by phase and baseline correction using the Bruker XWinNMR (V 3.5). The metabolites were identified and quantified using R package ASICS and statistical analysis was performed with Metaboanalyst 3.0.

Mitochondrial morphology calculations:

High resolution images of mitochondria stained with Tom20 were obtained using Zeiss 880 system, 63x oil objective and used for analysis. Approximately 40 cells were analyzed per condition with MiNA plugin for image j (10). It was obtained as per instructions provided (<https://imagej.net/MiNA_-_Mitochondrial_Network_Analysis>) and used to calculate mitochondrial morphology. The plots were generated in R and statistical tests were performed in PRISM^TM^.

### Expansion microscopy

Cells were cultured at a density of 4x10^4^ on glass coverslips, washed twice in PBS and fixed in 4% PFA in PBS. Following permeabilization in 0.15% Triton-X100 and blocking in 3% BSA, cells were probed with primary antibodies in the blocking buffer overnight at 4^o^C in a humidified chamber. The next day, samples were washed and incubated with ProExM compatible secondary antibodies: anti-mouse Alexa 488 and anti-rabbit Atto647N for 1hr in the dark and then washed before the sample preparation for the process of expansion. The ProExm was performed as described before(11). Briefly, immunostained cells were subjected to an anchoring treatment using 1:100 dilution of 10mg/ml stock solution of Acryloyl-X, SE (6-((acryloyl) amino)hexanoic acid, succinimidyl ester overnight at RT. The next day cells were encased in gelation solution (1x PBS, 2M NaCl, 8.625% (w/w) sodium acrylate, 2.5% (w/w) acrylamide, 0.15% (w/w) N,N’-methylenebisacrylamide, 0.2% (w/v) TEMED and 0.2% (w/v) APS) for 1hr at 37^o^C in a home-made chamber. Then the gel-encased coverslips were placed directly in the imaging glass bottom 6 well plate, immersed in digestion solution (Proteinase K, 50mM Tris pH 8.0, 1mM EDTA, 0.5% Triton X-100, 1M NaCl). Digestion was performed overnight in the dark, while mixing gently. The next morning gel pellets containing anchored and digested cells were washed multiple times in _dd_H_2_O to physically expand the samples, the Hoechst 3342 staining was performed during the last wash at 1.62uM concentration. The expanded gels were then covered with 1% low melt agarose in water to prevent sample movement during confocal imaging.

Confocal imaging was performed using Zeiss 880 system using 100x oil objective (1.4 NA) and AiryScan mode (where indicated). ZEISS Zen (blue edition) software was then used to deconvolute the 2D AiryScans for optimal signal to noise quality. Image analysis was performed using Fiji software (ImageJ).

CRISPR screen

CRISPR screen was performed using Brunello library acquired from Addgene. Briefly, Brunello library was transformed in electrocompetent cells and plasmids isolated using Qiagen maxi prep kits. The expanded library was prepared for illumina sequencing with requisite adaptors attached by PCR. Quality of sgRNA representation was confirmed prior to virus generation. On successful library expansion, HEK 293cells were prepared for virus production. Expanded Brunello library was transfected into HEK 293 cells using lipofectamine 2000 transfection reagent according to manufacturer’s protocol and virus containing supernatant collected at 48 and 72 h post transfection. MCF10A-ARID1A WT and KO cells were cultured in 6 well plate and serial dilution of virus was added to the cells and 1 μg/ml puromycin was used for selection. Appropriate viral volume was per cell was calculated and used for the experiment with MCF10A-ARID1A WT and KO cells. A total of 80 million cells per condition were cultured (1000 cells/sgRNA) in 15 cm dishes, calculated amount of virus was added to the cells and selected for 2 week in 1 μg/ml puromycin containing media. Upon selection cells were treated with DMSO/Volasertib for nearly a month to allow for cell kill and recovery of surviving cells. The surviving cells were harvested and genomic DNA isolated using Qiagen genomic DNA isolation kit. Concentration of the isolated DNA measured using nandrop and used for performing PCR with illumina adaptor primers. The PCR products were run on agarose gel to verify the band size of expected length and then mixed in equimolar ratio’s for illumina sequencing. Post sequencing the data was aligned and analysed following established standard protocols.

Multiplexed quantitative immunohistochemistry (mIHC)

Multiplexed quantitative immunohistochemistry (mIHC) was performed on formalin-fixed paraffin embedded (FFPE) samples to assess protein expression using an Opal 7 Color Kit and imaged and analyzed using the Vectra 2 System (PerkinElmer Inc., Waltham, MA, USA). Details of creating a quantitative protein nuclear expression score (NES) have been described previously(12) and applied here to create a ARID1A protein loss assay.

**Supplementary Tables**

**Supplementary Table 1: Antibodies used in this study**

| **Antibody** | **Company/clone** | **dilution** |
| --- | --- | --- |
| γ-H2AX | Merck Millipore, clone JBW301 05-636 | 1:1000 |
| pChk1( Ser345) | Cell Signaling technology, cat#2341 | 1:1000 |
| Total Chk1 | Santa Cruz-SC 8408 | 1:1000 |
| cleaved PARP | Cell Signaling technology, cat#5625 | 1:1000 |
| cleaved Caspase3 | Cell Signaling technology, cat#9661 | 1:1000 |
| HSC70 | Santa Cruz, SC-7298 | 1:1000 |
| PLK1 | Cell Signaling technology, 208G4 | 1:1000 |
| ARID1A | Bethyl laboratories A301-041A | 1:1000 |
| Phoshpho histone H3 | Cell Signaling technology, #9701 | 1:100 |
| Alpha tubulin | Abcam, ab7291 | 1:1000 |
| ATP5B | Abcam ,ab14730 | 1:1000 |
| Tom20 | Santa Cruz , SC-17764 | 1:500 |
| LC3B | Cell Signaling technology, cat#3868 | 1:1000 |
| P62 | Santa Cruz ,SC-28359 | 1:1000 |
| PARK2 | Santa Cruz ,SC-13367 | 1:1000 |
| Phospho DRP1 | Cell Signaling technology, cat#4867 | 1:1000 |
| DRP1 | Cell Signaling technology, cat#5391 | 1:1000 |
| ß-actin | Sigma, A5316 | 1:1000 |
| Alpha tubulin | Abcam ,ab15246 | 1:1000 |
| Alexa 488 | Thermofisher Scientific, A32790 | 1:1000 |
| Alexa 647 | Thermofisher Scientific, A32787 | 1:1000 |

**Supplementary Table 2: qPCR primers used in this study**

| **Primer name** | **Sequence (5'-3')** |
| --- | --- |
| PGC1a_FP | GGCAGAAGGCAATTGAAGAG |
| PGC1a_RP | TCAAAACGGTCCCTCAGTTC |
| PPARG_FP | TCTCTCCGTAATGGAAGACC |
| PPARG_RP | GCATTATGAGACATCCCCAC |

**Supplementary Figures**

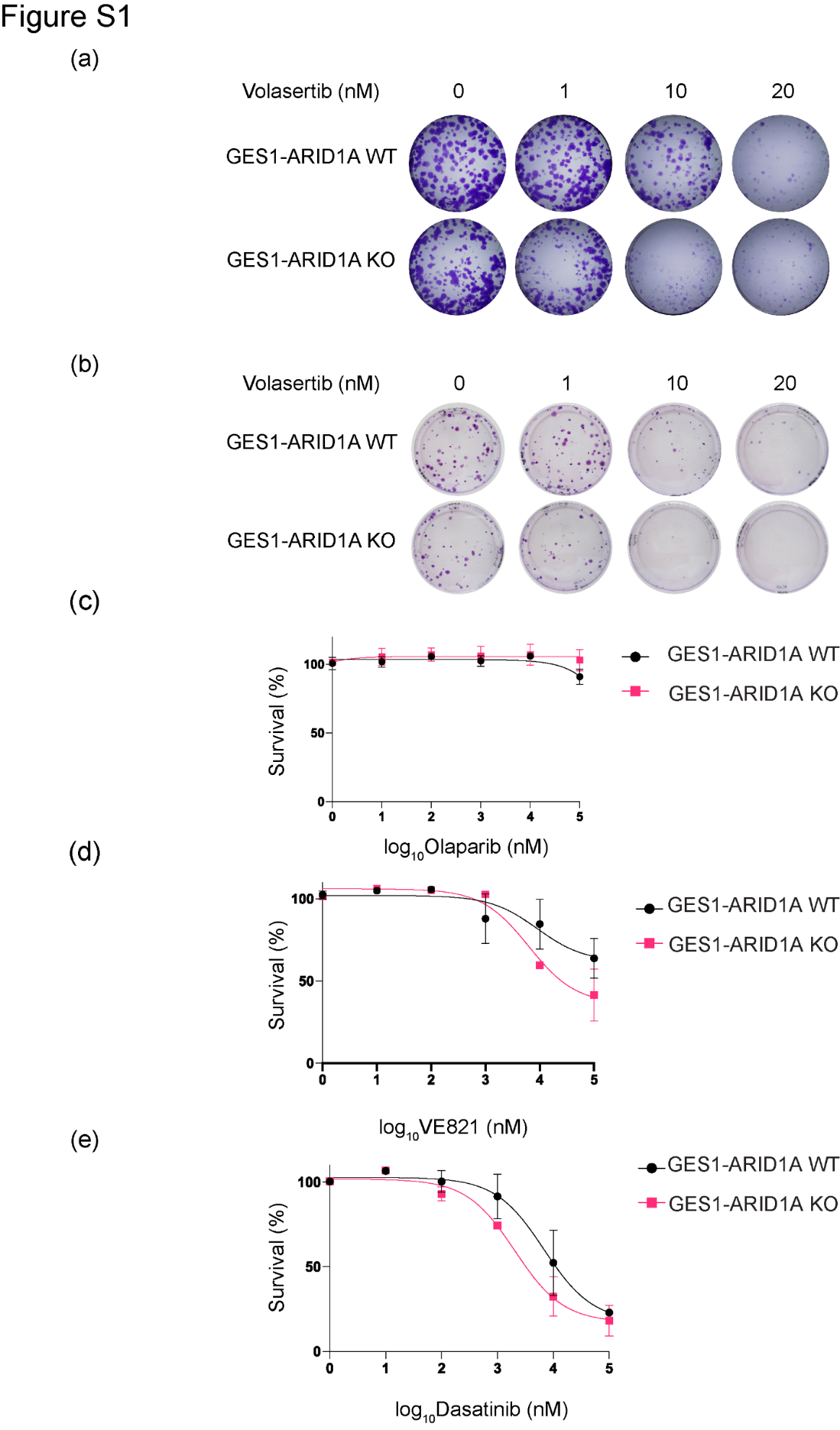

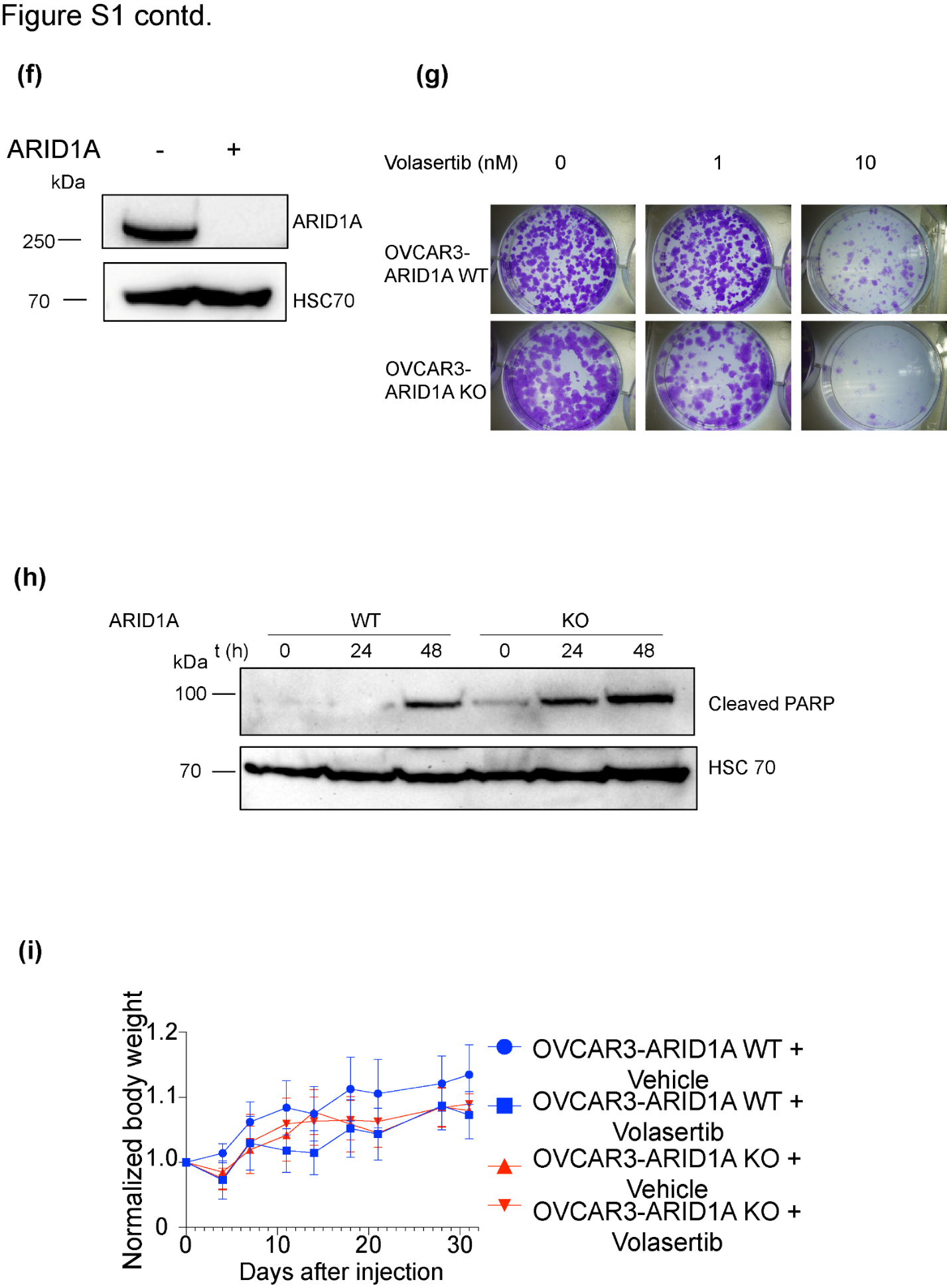

**Fig.S1:**

a) Representative images of a proliferation assay in GES1-ARID1A WT and KO cells treated with indicated concentrations of Volasertib and stained with crystal violet. Cells were plated at a dilution of 800 cells/well in a 6 well plate and treated with different concentrations of Volasertib for 10 days with fresh drug added every alternate day. Surviving cells were stained with crystal violet.
b) Representative images of colony formation assays quantitated in Fig. 1F, in GES1-ARID1A WT and KO cells treated with indicated concentrations of Volasertib.

Cell viability assays performed using cell titer blue in GES1-ARID1A WT and KO cells:

c) olaparib,
d) B) VE821, and
e) (C) Dasatinib. 1000 cells were plated on a 96 well plate, and treated with the indicated chemotherapeutics for 72 h. Cell titer blue was added to the cells and incubated at 37^o^C for 4 h. Fluorescence was measured with a plate reader and survival percentages were calculated. Graphs were generated in PRISM. Error bars represent SEM from 2 independent biological experiments.
f) Validation of OVCAR3-ARID1A KO cells by western blotting generated using CRISPR-Cas9 technology,
g) Cell proliferation assays showing synthetic lethality between ARID1A loss and Volasertib treatment in OVCAR3-ARID1A WT/KO cells,
h) Western blot showing increased PARP cleavage in OVCAR3-ARID1A KO cells in response to Volasertib treatment at various time points indicating higher apoptosis with time in OVCAR3-ARID1A KO cells relative to OVCAR3-ARID1A WT cells.
i) Plot showing the body mass of mice injected with OVCAR3-ARID1A WT and KO cells and treated with vehicular control and Volasertib.

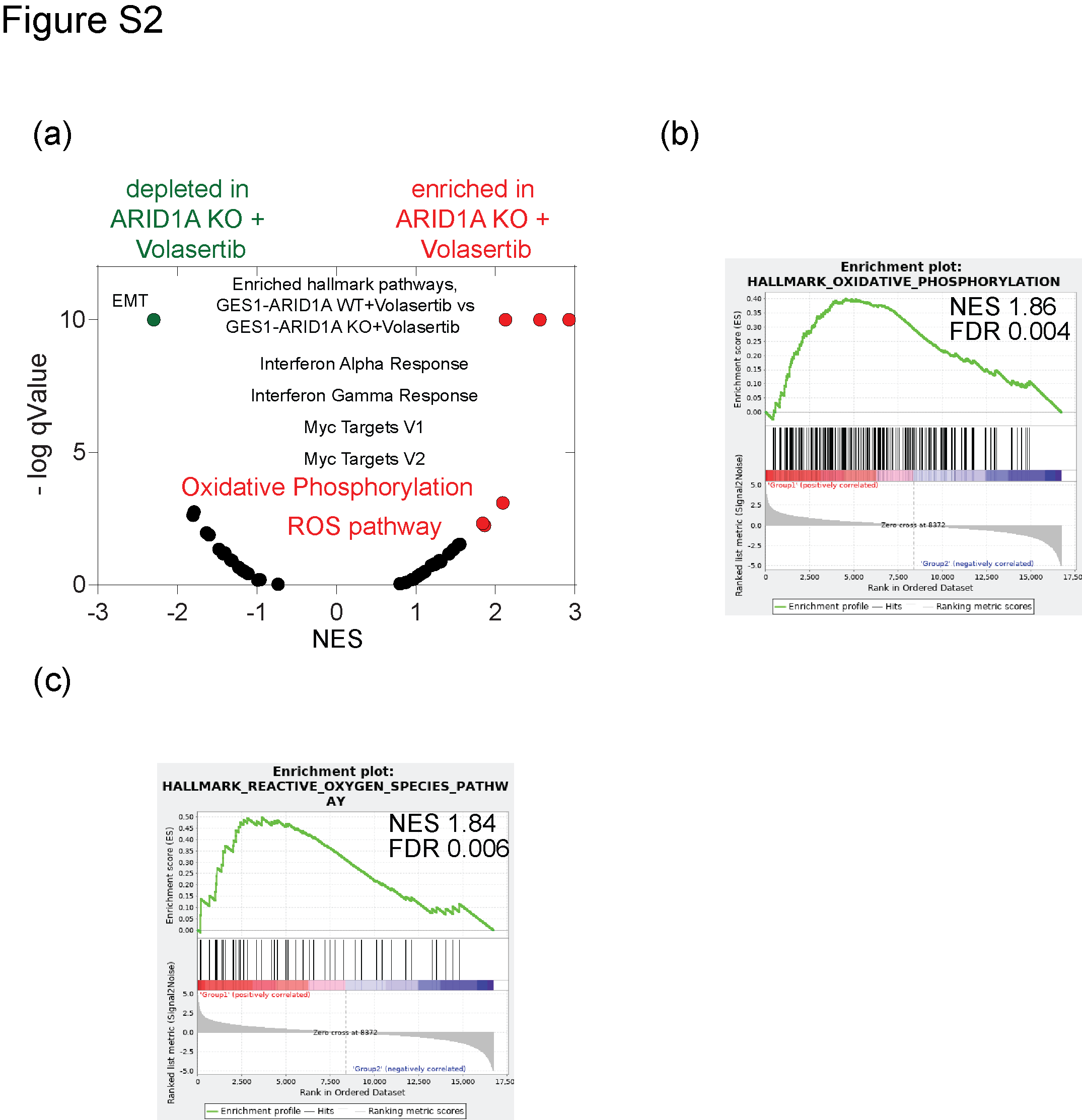

.

**Figure S2**

a) GSEA hallmark pathway enrichment analysis between volasertib treated GES1-ARID1A WT and KO cells showing enrichment in ROS pathway,
b) GSEA enrichment plot with normalized enrichment score (NES) for oxidative phosphorylation pathway between volasertib treated GES1- ARID1A WT and KO cells,
c) GSEA enrichment plot with normalized enrichment score (NES) for ROS pathway between volasertib treated GES1- ARID1A WT and KO cells.

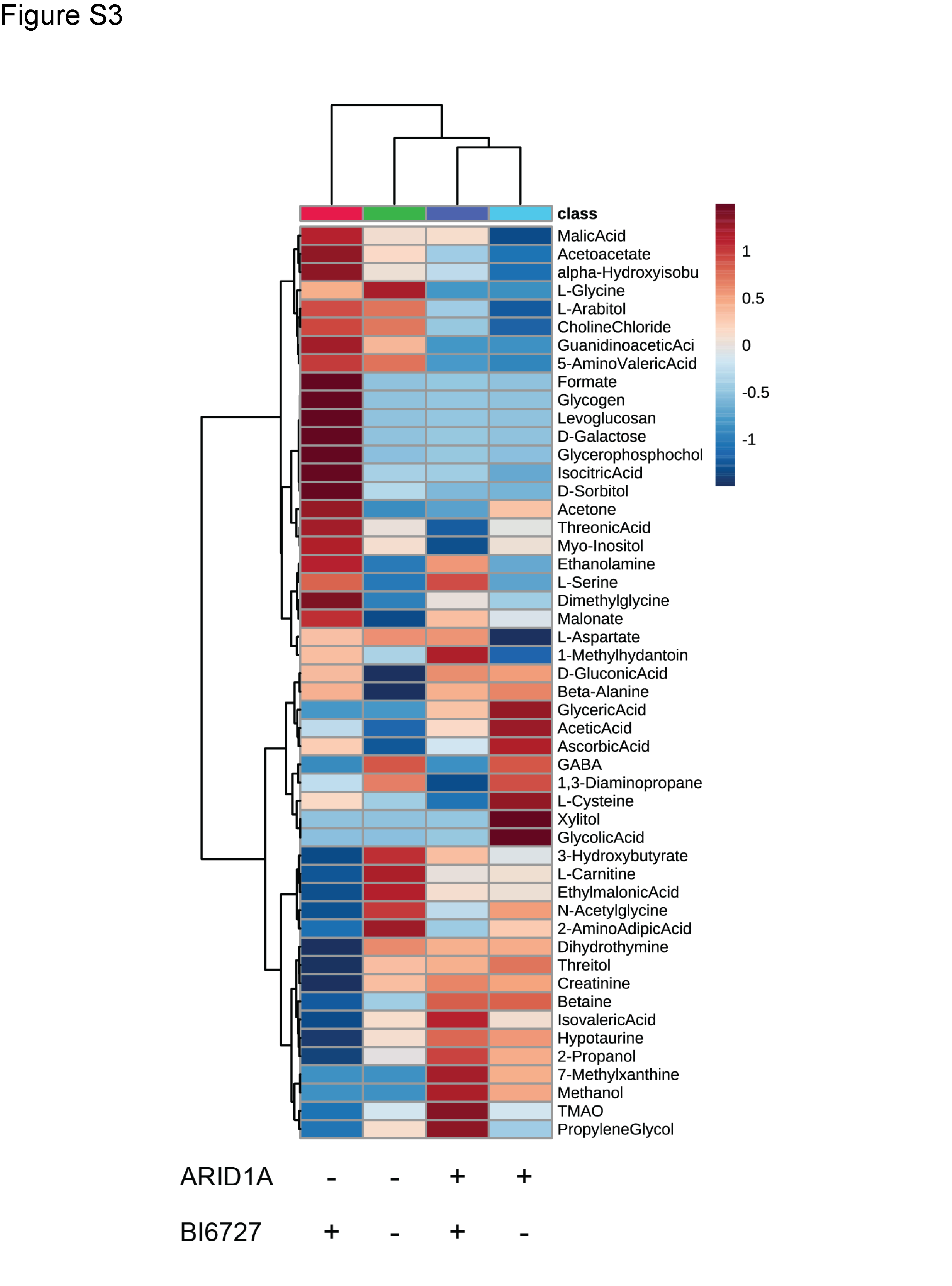

**Figure S3**: **1D NMR of ARID1A KO and WT cells in the presence of PLK1 inhibition**

Heat map of important metabolites up/down regulated in GES1-ARID1A WT and KO with and without volasertib treatment, as measured by 1D NMR metabolomics
